# Identification and Classification of Reverse Transcriptases in Bacterial Genomes and Metagenomes

**DOI:** 10.1101/2021.01.26.428298

**Authors:** Fatemeh Sharifi, Yuzhen Ye

**Author notes:** Corresponding author: Yuzhen Ye.

## Abstract

Reverse Transcriptases (RTs) are found in different systems including group II introns, Diversity Generating Retroelements (DGRs), retrons, CRISPR-Cas systems, and Abortive Infection (Abi) systems in prokaryotes. Different classes of RTs can play different roles, such as template switching and mobility in group II introns, spacer acquisition in CRISPR-Cas systems, mutagenic retrohoming in DGRs, programmed cell suicide in Abi systems, and recently discovered phage defense in retrons. While some classes of RTs have been studied extensively, others remain to be characterized. There is a lack of computational tools for identifying and characterizing various classes of RTs. In this study, we built a tool (called myRT) for identification and classification of prokaryotic RTs. In addition, our tool provides information about the genomic neighborhood of each RT, providing potential functional clues. We applied our tool to predict and classify RTs in all complete and draft bacterial genomes, and created a collection that can be used for exploration of putative RTs. Application of myRT to gut metagenomes showed that gut metagenomes encode proportionally more RTs related to DGRs, outnumbering retron-related RTs, as compared to the collection of reference genomes. MyRT is both available as a standalone software and also through a website.

## INTRODUCTION

Reverse Transcriptase (RT) is an enzyme that converts RNA into cDNA, and was discovered in 1970 in retroviruses [1]. A well-known RT in retroviruses is HIV-1 RT which integrates the HIV-1 virus RNA into the DNA of the host (human)[2]. All the retrotransposons (LTR and non-LTR) have RT genes [3]. Bacterial RTs were first found in retrons retroelements. Bacterial RTs can also be found in bacterial defense systems against phages, e.g. in CRISPR-Cas systems and abortive infection systems (AbiA, AbiK, Abi-P2)[4]. RTs are also found in Diversity Generating Retroelements (DGRs) that facilitate tropism switching in phages, and accelerate the evolution of bacteria and archaea [5]. Another important class of RTs can be found in mobile retroelements such as group II introns (GII/G2I) [6].

RTs involved in group II introns are the most abundant class of RTs in bacteria, and encode 57%-75% of the bacterial RTs [4, 6]. Bacterial group II introns are self-splicing mobile elements, each consisting of a catalytic RNA and an intron-encoded protein (IEP) within the RNA. The IEP contains a RT domain, X/thumb domain with maturase activity, along with DNA binding (D) and endonuclease (En) domains [4, 7]. Group II intron RTs and their template switching mechanisms are used in different gene and genome editing techniques including targetron and thermotargetron [8–10].

Retrons encode 12%-14% of the bacterial RTs and are the second frequent class of RTs [4, 6]. Although retrons were discovered three decades ago, their function remained unknown until recently that they were found to function as phage defense mechanisms [11, 12]. Bacterial retrons are non-LTR-retroelements that produce multicopy single-stranded DNAs (msDNAs). Most retrons consist of msr-msd sequence and a RT gene. Retrons can also encode toxin/antitoxin systems, which can be triggered or blocked by phage proteins [13, 14]. Retron RTs have been suggested as a tool for precise genome editing techniques (e.g. CRISPEY and SCRIBE) as retrons can produce msDNA, and edit the target sequences [6, 15].

RT genes are an essential component of Diversity Generating Retroelements (DGRs)[16]. DGRs are found in bacteria, bacteriophages, and intraterrestrial archaea and archaeal viruses [17]. DGRs are beneficial to the evolution and survival of their host; for instances, they can mediate tropism switching in Bordetella phage [18], mediate bacterial surface display [19], have a role in regulatory pathway tuning [20], and impact the underlying temperate phage-bacteria interactions in human gut microbiome [21, 22]. RTs in DGR systems are special, in the sense that they are error-prone. The RTs generate hyper-variable regions in specific target genes (tail fibre protein, receptors, etc.), through a process called mutagenic retrohoming in which a template region (TR) is reversed transcribed into mutagenized cDNA (A-to-N mutations), and is replaced with a region in the target gene which is similar to the TR region, and is called variable region (VR) [5]. Analyses of the target genes of DGR systems have shown that some pfam domains commonly seen in target genes include, but are not limited to DUF1566, FGE-sulfatase, and Fib succ major [23].

RT genes are found in some classes of CRISPR-Cas (the bacterial adaptive immune) systems including several subsets of type III (III-A, III-B, III-C, and III-D) and type VI-A CRISPR-Cas systems that can acquire spacers directly from both DNA and RNA [24]. CRISPR-Cas RTs are believed to have been emerged from multiple occasions: CRISPR-Cas RTs in archaea (Methanomicrobia) branch from class F of group II introns, CRISPR-Cas RT (ABX04564.1) in *Herpetosiphon aurantiacus* falls into the group II intron clade, CRISPR-Cas RT in *Haliscomenobacter hydrossis* is related to retron RTs [25]. CRISPR-Cas RT in *Haemophilus haemolyticus* is related to Abi-P2 RTs, and is associated with type I-C CRISPR-Cas systems [26]. *Streptomyces* spp. has several CRISPR-Cas RTs associated with type I-E CRISPR systems. As RNA activity is not common in type I CRISPR systems, experimental study of these CRISPR-Cas RTs may result in interesting findings [27].

RTs are also found in three types of abortive bacteriophage infection (Abi) systems including AbiA, AbiK and Abi-P2 [4, 28, 29]. Abi systems are a type of bacterial defense mechanisms that can lead to programmed death of a phage-infected cell, in order to protect the surrounding cell(s), and are often encoded by phages (e.g. P2 prophage of *E. coli*), and plasmids of bacterial genomes such as *Lactococcus lactis* [30]. *L. lactis* has more than 22 different abortive infection systems (AbiA to AbiV) [31], among which, only two of them (AbiA and Abik) have a RT domain. Although AbiA and AbiK only share 23% identity, they both can stop phage P335 maturation by means of un-templated synthesis of a DNA covalently bonded to the reverse transcriptase domain in order to target the Rad52-like phage recombinases [32]. C-terminal HEPN domain of AbiA (HEPN AbiA CTD), which may promote cell suicide through RNase activity, is fused to RT encoded by a gene found in an operon containing other genes including restriction modification system (RM system) [31].

Uncharacterized RTs are encoded by conserved ORFs in bacterial genomes, but their exact function and classification are unknown. Nevertheless, a few studies have suggested groupings of these RTs based on different criteria such as previously published data, alignment of RT motifs (sequence conservation of the RT motifs), and similarity of their fused protein motifs [4, 6, 33]. The genomic neighborhood of these RTs can also provide us with information about the functions of these RTs: for instance, RTs of unknown classes 1 and 5 are fused with nitrilase motif in the C-terminal, RTs of unknown class 3 and class 8 tend to co-occur, unknown class 4 RTs have a fimbrial domain, and unknown class 10 of RTs have fused primase domain, suggesting a concerted priming and reverse by the protein that harbors these two domains [6]. Despite the grouping, a few RTs remain unclassified as they don’t seem to have any close relationship with the other RTs in the collected dataset of RTs [6]. A recent study, discovered that six classes of unknown RTs, including unknown class 3 and unknown class 8 are part of the defense systems against dsDNA phages [34]

Due to the importance and applicability of bacterial reverse transcriptases, there are tools and databases that have been developed for individual classes of RTs, or genetic elements that contain the RTs. There is a database of group II introns (http://webapps2.ucalgary.ca/groupii/) [35, 36]. MyDGR is a tool that we developed for identification of DGR systems and their associated RTs [5]. However, a tool for characterization and classification of RTs remains lacking. We provide here the first pipeline for prediction of bacterial RTs and their classes, accompanied by genomic neighborhood information and visualizations. Furthermore, our pre-computed collection of putative RTs in all complete and bacterial genomes is easily accessible through myRT web-server.

## MATERIALS AND METHODS

### Collection of the RT dataset

A dataset of 1,718 non redundant RTs was collected from different sources. CRISPR-Cas associated RTs were collected from [26], Bacterial group II intron RTs were extracted from groupii [35]. The DGR RTs dataset including 421 non-redundant RTs were previously collected as part of our research on DGRs systems [5], and come from multiple sources [23, 37–40]. We also downloaded nine AbiA (abortive AbiA) representatives from CDD [41]. These 4 RT datasets were combined with a comprehensive dataset of RTs collected by [4]. As some of these datasets overlap, redundant RTs were removed (using cd-hit [42] cutoff value of 1). This integrated dataset contains RTs from group II introns (GII), CRISPR-Cas, DGRs, retrons, G2L4, AbiA, AbiK, AbiP2, UG1-10, UN1-5, and unclassified (UNC) RTs. This classification is mostly based on [4]; UG stands for unknown groups and UN stands for unclassified elements. We didn’t include unknown group 11, as the only element in this group is identical to a gene that is classified as CRISPR-Cas RT in [26]. Moreover, 2 out of 3 members of unknown group 11 in [6] are identical to RTs from retrons class in [4]. There remains one member (ACS60285.1) which also seems to belong to retrons. Therefore, there are no models for UG11 RTs in our study. Apparently Unknown group 10 class in [6] is different from unknown group 10 in [4], and contains an appended Primase (AE Prim S like) domain. And, group II like 3 (G2L3) class from [6] is UG10 (G2L3) class in [4]. As both of these classes seemed important, we renamed the one with Primase domain to unknown group 12 (UG12), and searched for UG12 RTs by searching for RTs with an appended Primase domain, and selected five of them manually to build the model for this class. We used a phylogenetic tree to verify that these RTs group together and fall in one clade (see Results).

### Construction of class-specific HMMs of RVT 1 domain for RT prediction and classification

Since all RT sequences contain the RVT 1 domain (Pfam ID: PF00078), we used the RVT 1 sequences to build class specific HMMs for RTs of different classes, which can then be used for identification and classification of RTs in genomes and metagenomes. For identification of RVT 1 domain in RT sequences, we used hmmscan (hmmer-3.2) [43] search against the Pfam-A model (PF00078) [44], and further validated the prediction using CD-search [41] and manual check. Hits of low significance or with a short length were manually checked.

We used Muscle (v3.8.31) [45] to align the extracted RVT 1 domains in the RT sequences, and then used FastTree2 (v3.8.31) [46] to build a phylogenetic tree of all the RT sequences, using bootstrap value of 100. By examining the phylogenetic tree, in combination with genomic context analysis, we confirmed the grouping of the RT sequences in the different classes, and for a small number of cases, re-assigned their classes (see Results). We also added a few small classes of RTs, including CRISPR-like. In total, all RTs can be grouped into 34 classes.

Extracted RVT 1 domains for each class were aligned separately using Muscle, and after re-formatting the alignments from fasta to stockholm, we used hmmbuild [47] to obtain hmm models for each class. Then, all of these hmm models (for different classes) were combined into one model (RVT-All.hmm).

### MyRT for identification and characterization of RTs in genomes and metagenomes

We developed myRT for identification of RTs in genomes and metagenomes. MyRT is based on similarity search against the class-specific RT HMMs, facilitated with phylogenetic analysis by pplacer for the cases when no clear classes can be inferred based on similarity search. First, FragGeneScan (version 1.31) [48] is used to quickly predict the protein coding genes in the input genome (or metagenome); however, if prediction of protein coding genes (given in a gff file) is available, protein sequences will be generated based on the input gff file instead. Next, our pipeline uses predicted protein sequences to find all the RVT 1 domains in the input genome in two steps (see Figure 1):

**Figure 1:**
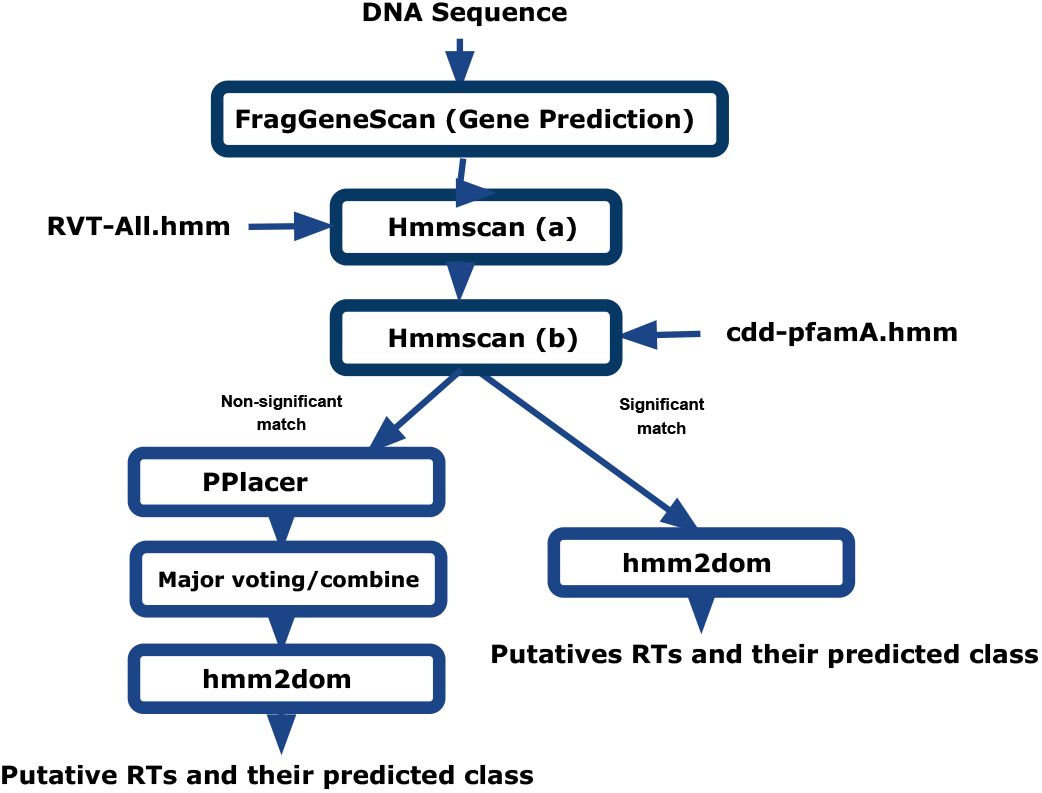
Flowchart of prediction of RTs and their class

a. Identification of initial putative RT proteins using a focused search of RT domains in all proteins. In this step, hmmscan is used to search all predicted proteins against RVT-All.hmm we created that contains only HMMs of class-specific RTs, with e-value of 10^*−*3^(-E 0.0001 –domE 0.0001) as the threshold. This step is very fast, as RVT-All.hmm only contains 34 sub-models. Proteins that are predicted to contain RT domains are considered candidates. Since RVT-All.hmm only contains RT domains, some of the identified candidates are likely to be false positives and need to be filtered out by the following step.
b. Refinement of RT protein candidates by expanded search of domains in the RT candidates. In this step, hmmscan is applied to search candidate RT proteins against HMMs of a large collection of domains (cdd-pfamA.hmm, which contains a total of 59,083 CDD and Pfam-A domains). Candidates that don’t contain a RT related domain (i.e., RVT 1, group II RT mat, RT G2 intron, etc) are considered false positives and are excluded from further analysis. This step is crucial for filtering the false positives (e.g. genes containing DNA binding domains), and the combination of this step with the previous step provides a fast test with a high precision and recall.

Classification of putative RT proteins is based on the above hmmscan search results, and in some cases an extra step of phylogenetic placement of the proteins in the tree of RTs. Given a putative RT protein, we consider that its RT class can be confidently assigned, if only one class of RT domain is found by hmmscan, or the top hit has a significantly lower E-value than the rest (i.e., E-value of the second hit is 10^5^ folds higher than the E-value of the top hit). In the cases when a class cannot be assigned, myRT keeps the top three hits, and relies on an extra step based on the placement of the putative RT protein in the phylogenetic tree of known RTs for final assignment of the class for the putative RT protein. The phylogenetic tree of known RTs was inferred by FastTree2 (with a bootstrap value of 100) using multiple alignment of RVT 1 domain sequences as the input. The reference tree (compatible with pplacer) was compiled using Taxtastic. To place a putative RT protein in the phylogenetic tree, first hmmalign is used to combine the putative RT with the reference hmm model. Then pplacer [49] is used to place the query sequence on the tree, and Treeio (v1.10.0) [50] and castor [51] R packages are used to parse the pplacer result. If the putative RT is placed on a leaf node, then the putative RT is assigned the class of the leaf node; otherwise if at least 90% of the leaf nodes in the subtree rooted at the putative RT share the same class, this class will be assigned to the putative RT. The confidence of the prediction at this step will be determined based on like weight ratio reported by pplacer, unless pplacer suggests several placements with similar like weight ratio, where the difference between second like weight ratio and first one is less than 0.25, in which we will report the result with a confidence value of 0. Finally, if the predicted class based on phylogenetic placement is consistent with hmmscan results (i.e., the prediction is among the top three hmmscan hits), this class will be selected as the final predicted class for the putative RT; otherwise, myRT reports all possible classes. We chose the parameters empirically.

### Genomic neighborhood analysis for putative RTs

To provide genomic neighborhood information of putative RTs, myRT examines up to four neighboring genes for each putative RT (up to two genes downstream, and up to two genes upstream of the RT, with a maximum intergenic region of 2kb). The neighboring genes together with the putative RT proteins will be searched against cdd-pfamA.hmm (using hmmscan and a maximum e-value of 10^*−*3^) to annotate the proteins encoded by these genes. The results can be used to infer domains that are frequently fused to the RVT 1 domains of the putative RT proteins, and the frequent domains encoded by the neighboring genes of the RT gene. We note when the predicted class of RT is CRISPR, but the putative RT has no Cas genes in its flanking genes. In this case, myRT will re-assign the class as CRISPR-like2 (second class of CRISPR-like RTs). The web version of myRT provides visualization of the prediction of putative RTs along with their genomic neighborhood.

### Genomes and metagenomes

We applied myRT to predict putative RTs in reference genomes (including complete and incomplete) and selected metagnenomes. Reference genomes were downloaded from the NCBI ftp website as of 10/22/2020. For complete genomes, we used NCBI’s prediction of putative coding genes, whereas for draft genomes, we used FragGeneScan [48] to predict protein coding genes. For metagenomes, the reads were trimmed using Trimmomatic [52], and paired reads were assembled using MetasSPAdes [48]. FragGeneScan was then used to predict putative coding genes from the assemblies of the metagenomes.

## RESULTS

### Reclassification of some RTs, expansion of rare RT classes and addition of new classes

We improved the collection of RT sequences and their models from three different aspects: reclassifying some RT sequences that were likely misclassified; adding more sequences for rare RT classes for model construction, and adding new RT classes.

We first applied the class-specific HMM models to predict and assign classes to the sequences in the initial training dataset. More than 99% of predictions agreed with the old classification. The rest could be either errors in the old classifications or misclassifications introduced by our method myRT. We analyzed these cases further, combining their sequential, genomic neighborhood, and phylogenetic information. Further we used CRISPRone [53], myDGR [5], and groupii [35, 36] to confirm RTs involved in CRISPR-cas, DGR, and group II introns, respectively. We excluded three sequences that don’t contain RVT 1 domain, including YP002455118.1 (WP000385107.1), KQB14190.1 and WP009625650.1. In addition, we were able to revise the classification for a total of 16 RTs summarized in Table 1. For example, sequence CBL40120.1 was previously labeled as Unclassified RT, however, it was predicted to be DGR RT by myRT, which was also confirmed by myDGR, a tool for DGR prediction. ZP 01872295.1, previously labeled as UG11, is identical to EDM23124.1 which is correctly labeled as CRISPR RT in [25]. EGP13976.1 (AEB93977.1) is one of the 155 DGR RTs identified by [38], but it is the only one (out of 155) that is not part of a DGR system; we regrouped it as an intron-RT with appended GIIM domain.

**Table 1:** Re-classification of 16 of the previously labeled RTs.

| Accession number | Old classification | New classification | Co-occurring domain(s) |
| --- | --- | --- | --- |
| CBL40120.1 | UNC | DGRs | Phage_XkdX |
| ZP_01872295.1 (EDM23124.1) | UG11 | CRISPR-Cas | *cas1* |
| EGP13976.1 (AEB93977.1) | DGRs | Group II introns | GIIM<sup>c</sup> |
| YP_001397265.1 (EDK35894.1) | DGRs | Group II introns | Intron_maturas2<sup>c</sup> |
| AFZ16538.1 (WP_015180701.1) | UNC | AbiA | MazF, HTH_XRE |
| NP_442332.1 | UG3 | UG7 |  |
| AFY59940.1 | UG3 | UG7 |  |
| AGA07305.1 | UG6 | UN3 |  |
| EPZ72367.1 | UNC | UN4 |  |
| AGI67543.1 | Group II like 3<sup>a</sup> | UG3 | UG8 |
| AEJ99900.1 | Group II like 4<sup>a</sup> | UG4 | FimD, FimA |
| AGK68212.1 (YP_005196274.1) | UG11<sup>a</sup> | Retrons |  |
| CCF10237.1 (EQR96236) | UG11<sup>a</sup> | Retrons |  |
| WP_007781002.1 (EJL44959.1) | CRISPR-Cas<sup>b</sup> | DGRs | Avd_like |
| WP_015462025.1 | CRISPR-Cas<sup>b</sup> | UG10 | SLATT_5 |
| WP_036415250.1 | CRISPR-Cas<sup>b</sup> | UG10 | SLATT_5 |

<sup>a</sup> These entries are from [6]. <sup>b</sup> These labels are based on [25]. <sup>c</sup> These domains are encoded by the same gene that encodes the RT domain.

We improved the hmm models for the classes with few representatives. We applied myRT to find putative RTs in 20,036 complete prokaryotic genomes, and extracted new RTs that belong to classes with few representatives (for UN1 we also selected one RT from draft genomes). Then, we verified the accuracy of these new classifications by adding the RVT 1 motif sequence of these putative RTs to the phylogenetic tree of RTs, to make sure that they fall in the right clade. After adding these newly classified RTs to the training data, we rebuilt the hmm models for these rare classes.

We added a few new classes of RTs based on a combination of sequential, phylogenetic tree and genomic neighborhood analyses. The first new class is RVT-CRISPR-like, and its model was built using 13 RT sequences that are similar to RVT-CRISPR but are not found in the CRISPR-Cas loci; and these sequences are clustered together in a branch in the phylogenetic tree (see Figure 2). We also proposed three new UN classes, RVT-UN6, RVT-UN7 and RVT-UN8. The final RVT-All.hmm contains 34 HMM models built from a total of 1513 RVT 1 sequences. Number of representatives of each class, and multiple alignment of RVT 1 motif sequences of each class is available at [myRT-Alignments](https://omics.informatics.indiana.edu/myRT/alignments.php). See Figure 3 for a barplot of representatives in our final models compared with the previously labeled RTs.

**Figure 2:**
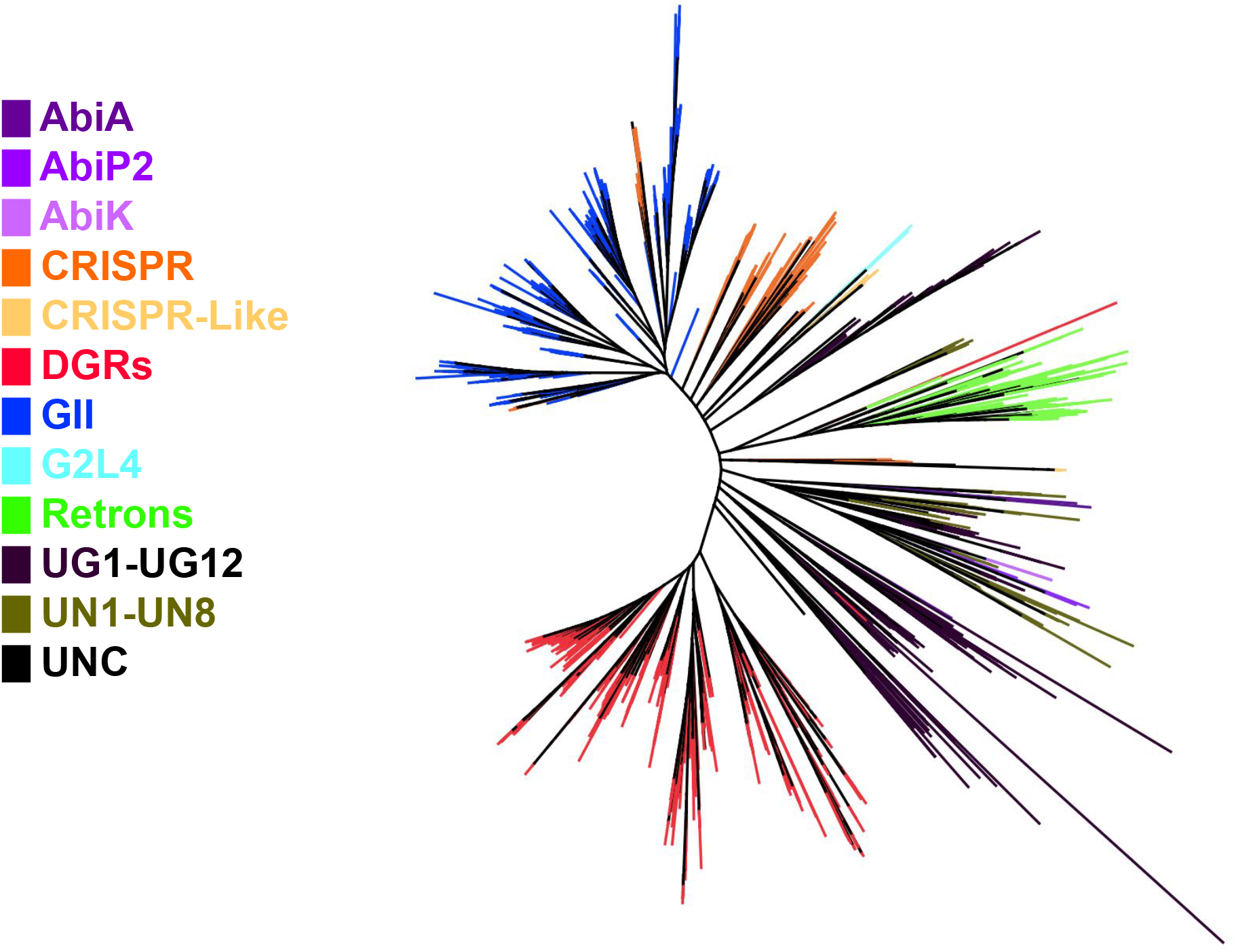
Phylogenetic tree of different RTs. The tree was inferred using the alignment of RVT 1 domains, and visualized using Archaeopteryx [54]. The branches are colored according to the RT classes.

**Figure 3:**
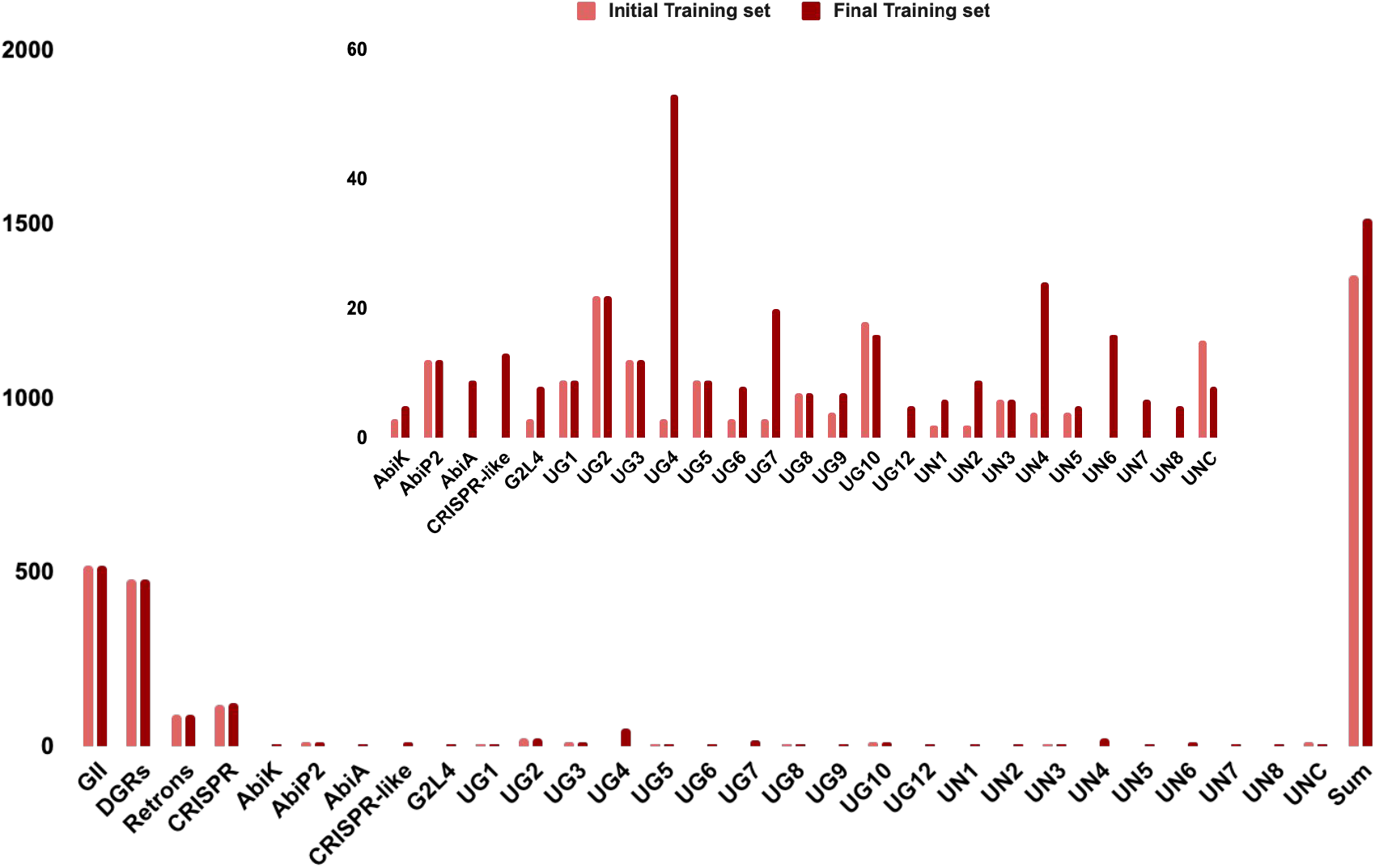
Barplots of the number of RTs in each class in our initial training set versus our final training set

### Evaluation of myRT using three independent collections of RTs

We applied myRT to three independent collections of RTs for evaluation. All results showed that myRT gave accurate classification of RTs. Supplementary Tables S8-10 provide access of myRT results for these three collections.

The first collection contains CRISPR-Cas reverse transcriptases (branch 1 - branch 10) from [27] that were not used in building the hmm models. The last column of Table 2 shows the identities of each RT to the RVT 1 motif sequences used in building RVT-All.hmm. KKO19091.1 (LAQJ01000220.1_7006_7917_-) despite being a CRISPR-RT shares 39% sequence identity with RVT 1 domain of AGB41082.1 (GII RT), and shares 37% sequence identity with WP 012599795.1 (CRISPR-Cas RT), and was correctly predicted as CRISPR-Cas RT by our pipeline solely based on hmmscan results. GAN31766.1 (RT#1 in BAFN01000001.1) shares 71% identity with CAJ74578.1, and based on CRISPRone results seems to be adjacent to a *cas4* gene, whereas CAJ74578.1 which is adjacent to a *cas1* gene. Our pipeline did not recognize the *cas4* gene, only reported a gene encoding GxxExxY appended domain, and thus labeled this RT as CRISPR-like2. Last three RTs in Table 2 are associated with type I-E CRISPR-systems, have an adjacent *cas3* gene, and an adjacent gene encoding AbiEii domain, which is not a usual genomic neighborhood for CRISPR-Cas RTs. We do not have any models for these unusual CRISPR-Cas RTs. Even though AJKO01000007.1 (*Streptococcus oralis* SK10) has a type I CRISPR system, the putative RT of this genome (EIC80228.1; AJKO01000007.1_124518_125894_-) is not located in the CRISPR loci, and is predicted as UG2 RT by our pipeline. Based on this comparison, our pipeline can predict a CRISPR RT as CRISPR/CRISPR-like RT with an accuracy of 96%.

**Table 2:** Evaluation of myRT on the Silas 2019 collection of CRISPR-Cas RTs [27].

| Genome | RT coordinates # | myRT prediction | Identity%& |
| --- | --- | --- | --- |
| ASPN01000006.1 | ASPN01000006.1_20247_22901_+ | CRISPR | 34 |
| AQRP01000065.1 | AQRP01000065.1_13318_16137_+ | CRISPR | 34 |
| JXXW01000010.1 | JXXW01000010.1_71385_73319_+ | CRISPR | 59 |
| ASAJ01000015.1 | ASAJ01000015.1_125843_128860_- | CRISPR | 56 |
| KL370780.1 | KL370780.1_93557_95149_+ | CRISPR | 35 |
| JWIO01000025.1 | JWIO01000025.1_10124_11221_+ | CRISPR | 38 |
| BCQS01000026.1 | BCQS01000026.1_4694_5707_- | CRISPR | 40 |
| JH470356.1 | JH470356.1_27166_28113_+ | CRISPR | 77 |
| LLVU01000015.1 | LLVU01000015.1_5740_7752_- | CRISPR | 57 |
| LOAS01000028.1 | LOAS01000028.1_37027_38064_- | CRISPR | 48 |
| JYJP01000030.1 | JYJP01000030.1_21534_23426_+ | CRISPR | 74 |
| JH992891.1 | JH992891.1_86001_86924_- | CRISPR | 75 |
| ANNX01000115.1 | ANNX01000115.1_14418_15371_- | CRISPR | 76 |
| ALWD01000163.1 | ALWD01000163.1_24045_25013_+ | CRISPR | 85 |
| ANNX01000117.1 | ANNX01000117.1_21795_22772_- | CRISPR | 86 |
| JH992901.1 | JH992901.1_1016250_1017311_+ | CRISPR | 81 |
| JXCA01000005.1 | JXCA01000005.1_398870_399850_- | CRISPR | 82 |
| ALVY01000183.1 | ALVY01000183.1_27387_28304_+ | CRISPR | 69 |
| HE972669.1 | HE972669.1_45936_46913_- | CRISPR | 72 |
| ALWB01000016.1 | ALWB01000016.1_12151_14244_- | CRISPR | 64 |
| ASMA01000004.1 | ASMA01000004.1_21607_22458_+ | CRISPR | 46 |
| JQFA01000004.1 | JQFA01000004.1_1149670_1151637_- | CRISPR | 85 |
| AJLK01000155.1 | AJLK01000155.1_941_2959_- | CRISPR | 86 |
| ASZN01000033.1 | ASZN01000033.1_1862_3346_- | CRISPR | 52 |
| LAQJ01000220.1 | LAQJ01000220.1_7006_7917_- | CRISPR | 37 |
| KK211136.1 | KK211136.1_10156_11052_- | CRISPR | 62 |
| BAFN01000001.1 | BAFN01000001.1_294229_295173_+ | CRISPR-like2 | 71 |
| AJKO01000007.1 | AJKO01000007.1_124518_125894_- | UG2* | 34 (UG2) |
| DS570667.1 | DS570667.1_23830_25101_- | UNC* | 29 |
| CP007699.1 | CP007699.1_7617452_7618696_+ | CRISPR-like* | 38 (GII) |
| LAKD01000050.1 | LAKD01000050.1_46946_48151_- | CRISPR-like* | 25 |

\* These 4 RTs were referred as “RTs with unusual RT-associated CRISPR-Cas architectures” in [27]. #: Coordinates of the predicted RT genes are presented in the genome/contig\_start\_end\_strand format, which shows the start, end, and strand of the protein coding regions. & This column lists the highest sequence identity between the predicted RT and the RVT\_1 domains used to build the HMMs for the different classes of RTs.

We then tested our model on datasets from [6] (excluding unclassified RTs). To fairly assess myRT’s performance we excluded RTs that shared more than 87% identity with our RT collection. The majority of the retained ones share 30%-50% identity with our training set which is normal considering that all of them have conserved RT domains. The test set used for this evaluation, and the results can be found in Table 3. Most of the predictions matched with the groupings from this collection, except one G2L5 (GII-like-5 RT). This collection contains two G2L5 RTs (ZP 01854760.1 and ZP 01851752.1) from *Gimesia maris* DSM 8797. The two RTs share low (28%) sequence identity, and both seem to be located in transposons. ZP 01854760.1, which shares 33% sequence identity with YP 552148.1 (a GII RT), but lacks the GIIM domain, was correctly predicted by myRT as a GII RT with a DDE Tnp 1 (transposase) domain in its genomic neighborhood. However the other one ZP 01851753.1, which has HTH Tnp 1 (helixturn-helix) and Tra5 (transposase InsO and inactivated derivatives) in its flanking genes, was incorrectly predicted as CRISPR-like2 by myRT, resulting in 95% accuracy overall.

**Table 3:** Result of testing myRT on the Zimmerly 2015 collection of RTs [6].

| Accession number | Gene coordinates | Class | myRT prediction |
| --- | --- | --- | --- |
| CAA78293.1 | AHAX01000006.1_19205_20143_+ | Retrons | Retrons |
| AAM42896.1 | AE008922.1_4324395_4326089_+ | Retrons | Retrons |
| AFY43443.1 * | CP003548.1_3308170_3310188_- | CRISPR | CRISPR |
| ZP_01854760.1 | NZ_ABCE01000016.1_73333_76125_+ | G2L5 | GII |
| ZP_01851752.1 | ABCE01000001.1_117632_117919_- | G2L5 | CRISPR-like2 |
| EKP98429.1 | CP006690.1_1914333_1915748_- | UG2 | UG2 |
| CCC73043.1 | HE576794.1_939599_941005_+ | UG2 | UG2 |
| AEG09910.1 | CP002767.1_716105_717388_- | UG2 | UG2 |
| AEW72297.1 | CP002886.1_903224_904396_+ | UG3 | UG3 |
| ACD38707.1 | EU595736.1_18886_20160_+ | UG3 | UG3 |
| AHE72406.1 | CP006580.1_34213_36348_- | UG4 | UG4 |
| AGI74246.1 | CP003742.1_4621630_4622688_+ | UG4 | UG4 |
| CDI94624.1 | HG530068.1_6882792_6885812_+ | UG5 | UG5 |
| AHM47056.1 | CP007393.1_928229_931438_- | UG5 | UG5 |
| ACU61669.1 | CP001699.1_5262939_5266187_+ | UG6 | UG6 |
| AHL77683.1 | CP007441.1_3859031_3861010_- | UG8 | UG8 |
| AEN62840.1 | CP003026.1_82890_84845_- | UG8 | UG8 |
| AFG37103.1 | CP003282.1_1170484_1172610_- | UG8 | UG8 |
| ADI30165.1 | CP002056.1_2016134_2017783_- | UG9 | UG9 |
| AGA65410.1 # | CP002873.1_44531_46162_+ | UN5# | UN5 |

\* We note that AFY43443.1 was labeled as a G2L1/G2L2 RT in [6], but the authors reported that the RTs classified as G2L1 and G2L2 are associated with *cas1* genes of CRISPR/Cas loci. G2L1 and G2L2 were proposed originally in [33], and in Toro et al [4], they were renamed as CRISPR-RT because of their association with the CRISPR-Cas systems. # AGA65410.1: this RT was classified as UG14 in [6], which includes three UG14 RTs including AGA65419.1, BAL31358.1 and AEA43024.1; all these three sequences were classified as UN5 by myRT, which is consistent with Toro et al's classification, which classified BAL31358.1 as a UN5 RT [4].

Finally, we tested our pipeline on a dataset of 16 experimentally retrons with experimentally validated RTs [15], 12 of which were recently experimentally proved to function as anti-phage defense systems. Five of these RTs were already in our training set, yet the other 11 shared less than 61% sequence identity with our training data (RVT-All). MyRT was able to precisely predict all of them as retron RTs, providing an accuracy of 100% (see Table 4).

**Table 4:** Evaluation of myRT on the Simon 2019 collection of retron RTs [15], all of which were predicted as retron RT by myRT.

| Known Retron | Genome | RT coordinates | Identity%<sup>&</sup> |
| --- | --- | --- | --- |
| EC48<sup>#</sup> | *Escherichia Coli* DE147 | LFQP01000005.1_154506_155696_- | 50 |
| EC67<sup>#</sup> | *Escherichia coli* S10 | CP010229.1_4712073_4713833_- | 61 |
| EC73<sup>#</sup> | *Escherichia coli* M10 | CP010200.1_2393178_2394128_+ | 36 |
| Ec78<sup>#</sup> | *Escherichia coli* 102598 | JHRW01000018.1_27622_28557_+ | 48 |
| EC83<sup>#</sup> | *Escherichia coli* 05-2753 | CXYK01000012.1_74586_75524_+ | 47 |
| Mx65 | *Myxococcus xanthus* DSM 16526 | FNOH01000027.1_37959_39242_+ | 53 |
| Eco8<sup>#</sup> | *Escherichia coli* 200499 | CYGJ01000003.1_369367_370491_+ | 47 |
| Se72 | *Salmonella enterica*<sup>*</sup> | AMMS01000284.1_2640_3671_- | 48 |
| Vc137<sup>#</sup> | *Vibrio cholerae* 2012EL-1759 | JNEW01000012.1_609188_610135_+ | 49 |
| Vp96 | *Vibrio parahaemolyticus* S119 | AWJG01000250.1_32_1054_+ | 49 |
| YF79 | *Yersinia frederiksenii* ATCC 33641 | KN150731.1_1692670_1693602_- | 50 |

<sup>#</sup> These retrons function as anti-phage defense systems. <sup>\*</sup> *Salmonella enterica* enterica sv. Heidelberg 579083-10. <sup>&</sup> This column lists the highest sequence identity between the predicted RT and the RVT\_1 domains used to build the HMMs for the different classes of RTs.

The Ec48 retron system which has proved vigorous defense against *Siphoviridae, Myoviridae*, and *Podoviridae* phages, has an Abi-P2 RT in its genomic neighborhood, which shares 99% identity with CAJ43157.1, Abi-P2 RT of *Enterobacteria* phage P2-EC58. Two other genes in the genomic neighborhood of the Ec48 Retron system have predicted Q (portal vertex) and phage GPA (Bacteriophage replication gene A protein) domains.

### Putative RTs identified in bacterial genomes

We applied myRT to predict putative reverse transcriptase, alongside their class in all complete and draft bacterial genomes. In total, 8,251 out of 20,036 complete genomes, and 118,883 out of 262,497 draft genomes each contain at least one putative RT. This collection is easily accessible through our web-server. We note that for genomes with predicted RTs associated with DGRs or CRISPRs, DGR prediction (by myDGR) and CRISPR-Cas prediction (by CRISPRone) will also be provided. Figure 4 shows the distribution of each RT class in complete and draft bacterial genomes. Just as expected, group II intron is the most prevalent class of RT (≈ 66%), followed by retron (∼13.5%). Retrons were recently found to provide phage defense mechanisms, and myRT predicted a total of 3,102 and 34,259 retron RTs in the complete and draft genomes, respectively. These putative retron RTs will be useful for further study of the function and distribution of the retron RTs in bacterial genomes. MyRT results for these reference genomes and plasmids are available in Supplementary Table S5. Below we show several cases of myRT predictions for demonstration purposes.

**Figure 4:**
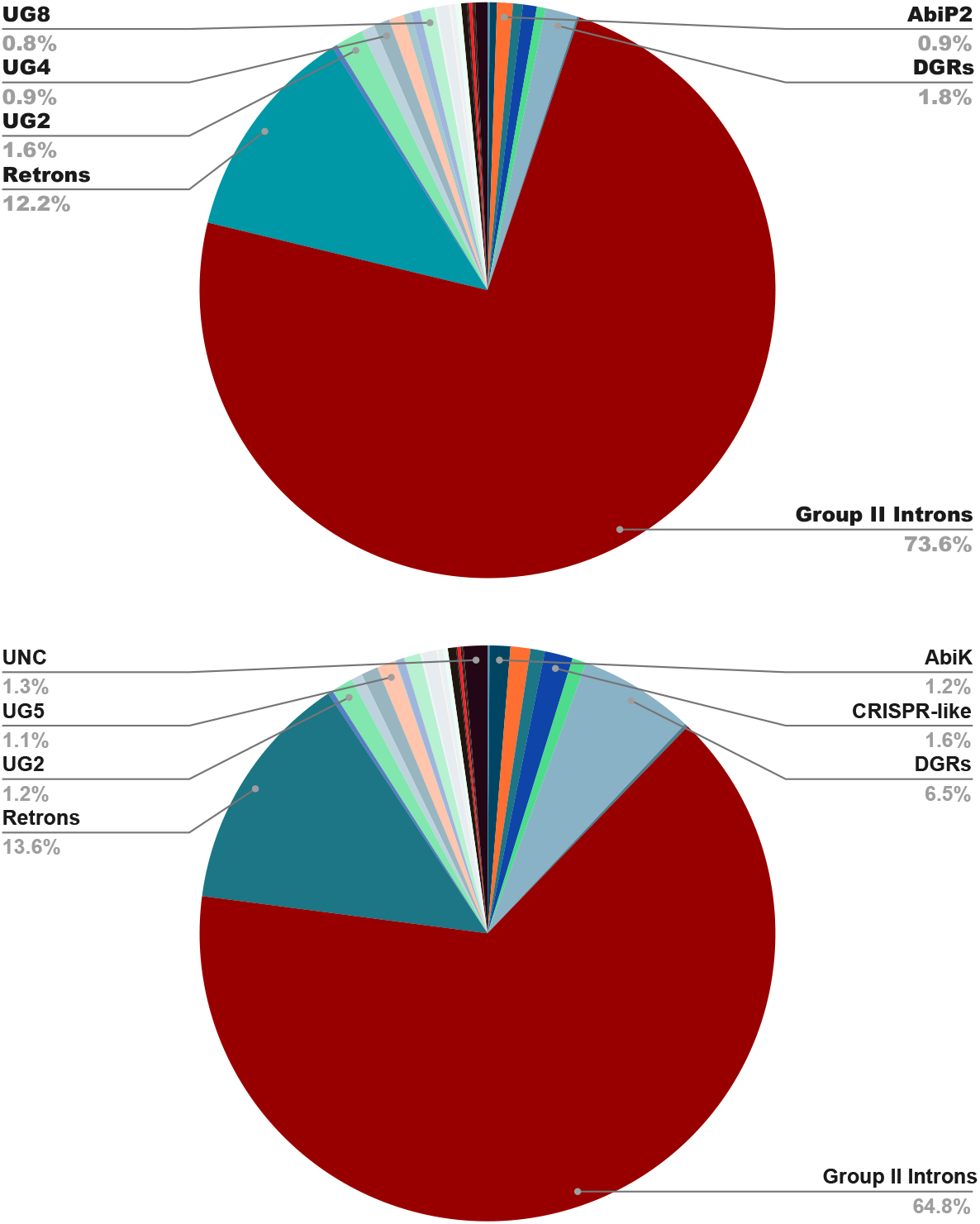
Distribution of different classes of RT in complete (top) and draft (bottom) bacterial genomes.

Figure 5 shows myRT predictions of two genomes. The first example is *Nostoc* sp. PCC 7120 and its plasmids. The two plasmids each contain a gene encoding group-II intron RT (not shown in the figure), and the genome encodes four classes of RT: DGR, CRISPR, retron, and group II intron as shown in Figure 5A. The second example is *Microcystis aeruginosa* NIES-843, which has seven RTs all related to group II introns. Six of the seven RTs are almost identical (sharing 97-99% identity), and they only share low identity (51%) with the seventh RT (which shares 65% identity with ACV02121.1, a group II intron RT in *Cyanothece* sp. PCC 8802). As seen in Figure 5B, six of these group II intron RTs have an appended McrA domain (5-methylcytosine-specific restriction endonuclease McrA).

**Figure 5:**
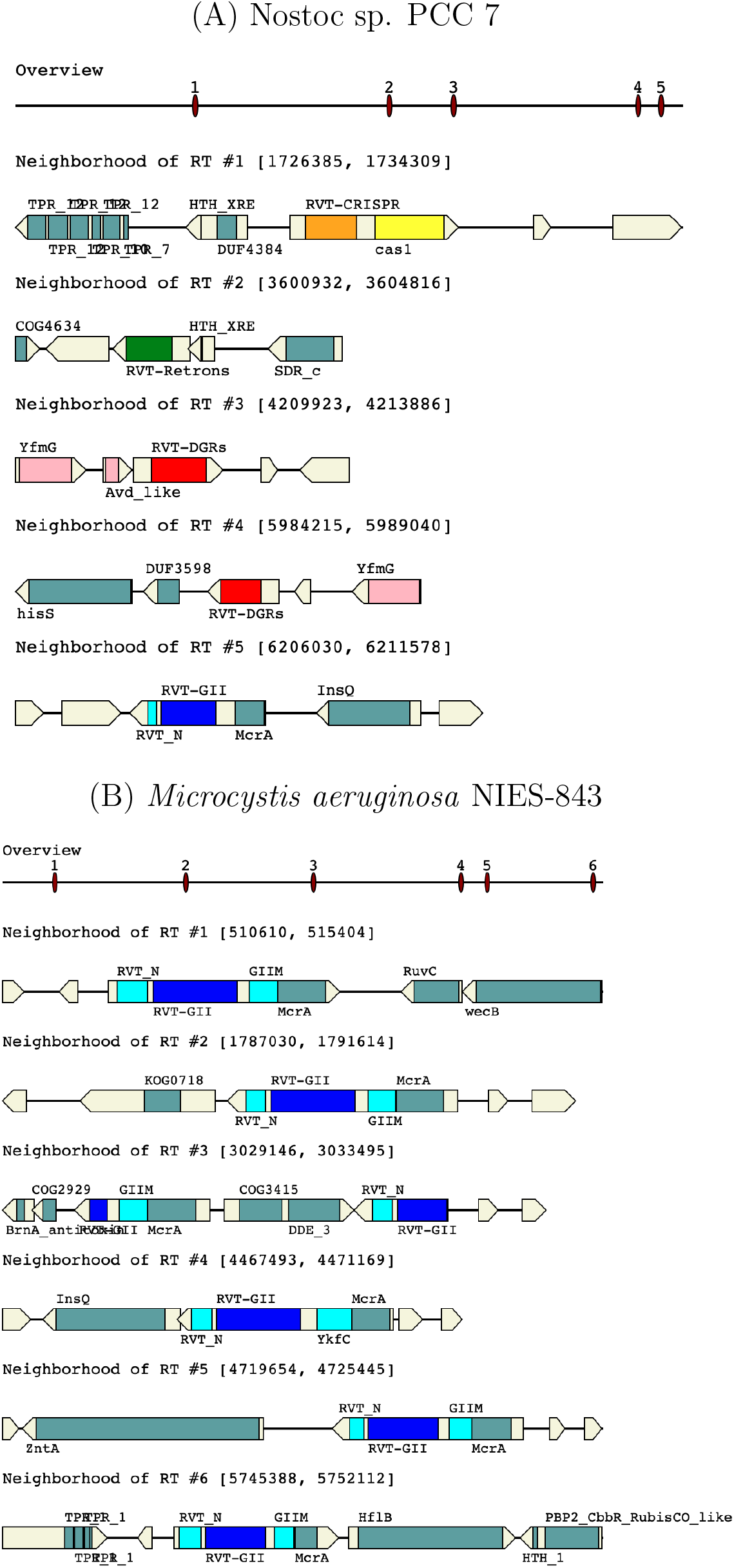
MyRT prediction results for (A) *Nostoc sp*. PCC 7120, and (B) *Microcystis aeruginosa* NIES-843. The “overview” view shows the locations of the predicted RT genes along the genome, and the zoom in views below each show one RT gene and its neighborhood. Arrows represent the genes, with the different regions encoding different domains in colored rectangles. All six RTs in *Microcystis aeruginosa* are group-II intron RTs (in blue), whereas *Nostoc sp*. contains RTs of different classes (in different colors).

Surprisingly, we observed that *Bacillus thuringiensis* YBT-1518 and its plasmids have a large number of group II intron RTs (74 GII RTs), and one RT with unknown function, which is similar to class of UG4 RTs, but doesn’t have Fimbrial domain in its genomic neighborhood. Out of these 74 RTs, 60 of them are identical to AHA69388.1 (RT #1), and 3 of them are identical to AHA69975.1 (RT #6). AHA69975.1 is adjacent to a potentially virulence gene with VirD4 (Type IV secretory pathway component) domain.

Application of myRT to the complete bacterial genomes resulted in the identification of 33 AbiA RTs. More than 60% of these AbiA RTs have an appended HEPN AbiA CTD. Examples of AbiA and AbiK, can be found in U17233.3 (*Lactococcus lactis* plasmid pTR2030), and in U35629.2 (*Lactococcus lactis* plasmid pSRQ800) respectively. Based on the genomic neighborhood, this AbiK seems to be part of a restriction-modification system. The third class of Abi RTs is Abi-P2, an example of Abi-P2 is found in *E. coli* 536. Some Abi-P2 RTs are located in a CRISPR-Cas loci, and may be associated with CRISPR-Cas systems as discussed earlier. Other examples of Abi RTs can be accessed through myRT-collection.

When applied myRT to predict putative RTs in complete genomes, about 97% of the putative RT had their class assigned solely based on hmmscan results. For the rest, phylogenetic information was used to assign a class to 80% of them. Among 25,487 predicted RTs in complete genomes, 172 RTs (less than 1%) remain unclassified, 46 of which share more than 56% identity with YP 003455357.1 (CBJ12261.1) unclassified RT in *Legionella longbeachae* NSW150. Table 5 contains some of the examples where phylogenetic information helped us to improve the classification of putative RTs. For instance ADE85032.1 was predicted as a CRISPR RT. Had we only used hmm models, it would be predicted as GII RT. This RT has a fused Cas1 domain, and its encoding gene has a *cas6* gene in the genomic neighborhood which indicates this RT is indeed related to a CRISPR-Cas system. The other set of examples include 10 UG3 RTs that otherwise would be classified as Unclassified RTs (e.g. UG3/DGRs/UG10), and we note that nine of these predicted UG3 have UG8 in their neighborhood (UG3 and UG8 tend to co-occur according to our training data and previous studies [6, 55].

**Table 5:**
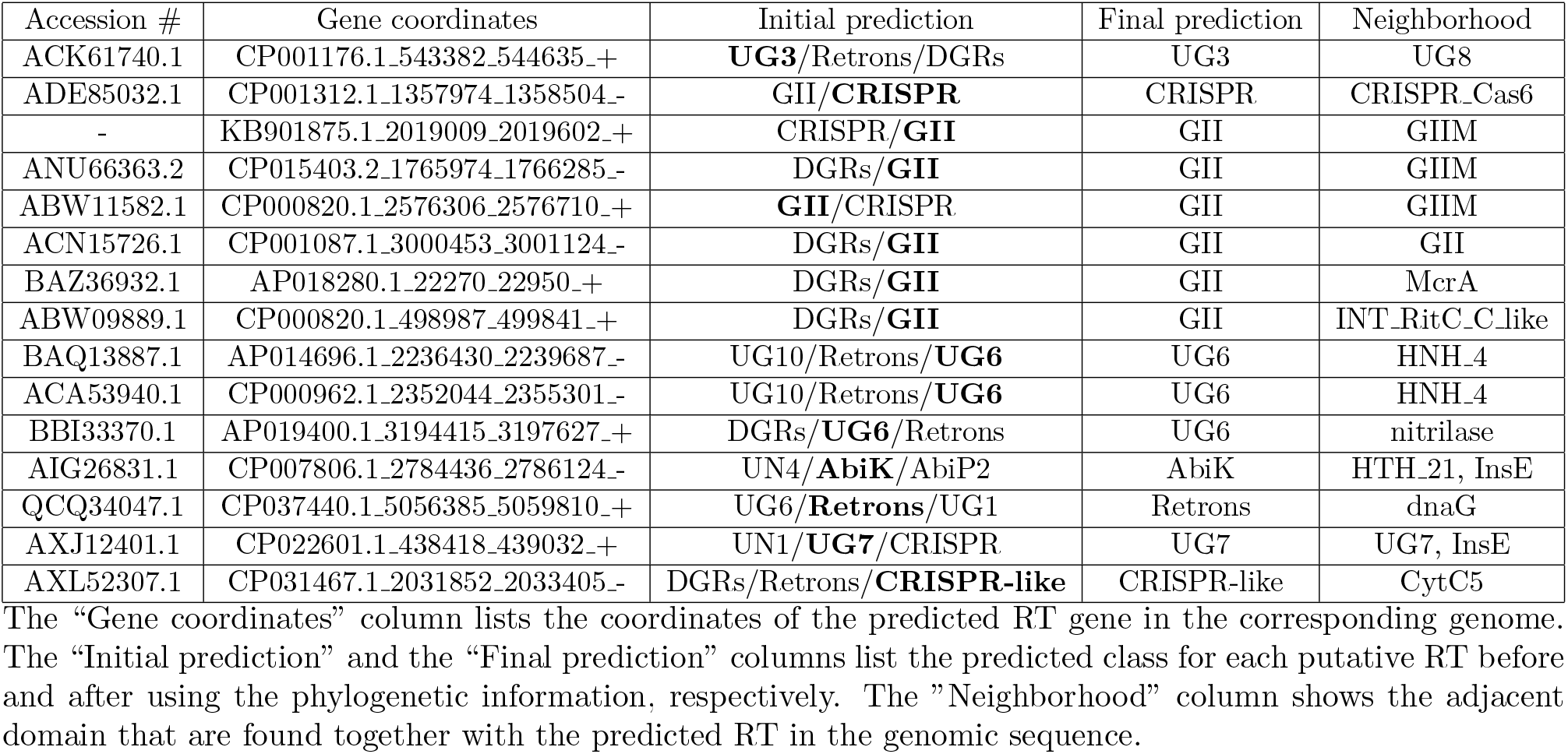
Potential improvements of RT classification by using phylogenetic information.

### Application of myRT to predict RTs in metagenomes

For demonstration purposes, we applied our pipeline to identify putative RTs in metagenomes. The first set contains four gut metagenomes (ERR248260-ERR248263) from fecal microbiota of human, chicken, cow, and pig from [56]. As seen in Figure 6, all gut metagenomes contain higher proportions of DGR-related RTs as compared to the reference bacterial genomes (see Figure 4), and the pig gut metagenome has the highest proportion of DGR RTs among all. The pig gut metagenome contains 1,333 putative RTs, including 654 GII RTs, 381 DGR RTs (352 after removing sequence redundancy by cd-hit [42] using sequence identify cutoff of 70%), 122 retron RTs and other classes of RTs. Out of 352 non-redundant DGR RTs in this dataset, 163 share less than 70% identity with the DGR-RTs from complete and draft bacterial genomes which contains 4,637 non-redundant DGR-RTs (cut-off value: 0.7). Using myDGR, we were able to identify 38, 15, 15, and 15 complete DGRs (a typical DGR system contains a RT gene, a template region TR, and a target gene containing the corresponding variable region VR) in the human, chicken, cow and pig gut metagenome, respectively, reflecting the fragmented nature of the metagenome assemblies (many of the contigs are very short). Supplementary Table S4 includes the links to myRT and myDGR predictions of these metagenomes.

**Figure 6:**
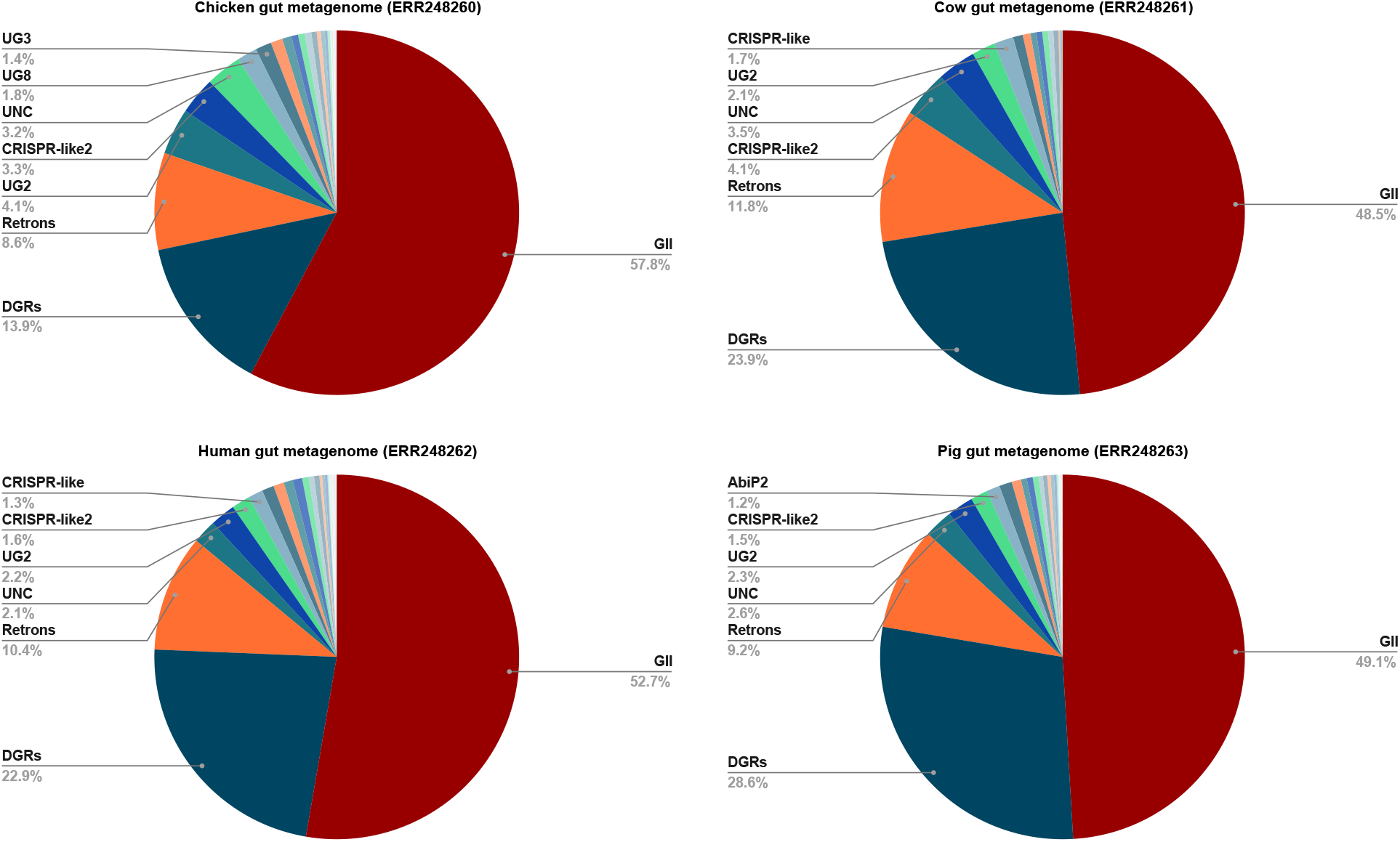
Distribution of RT classes in gut metagenomes of human, chicken, cow, and pig from [56]

To further investigate if pig gut metagenomes generally have a high proportion of DGR-RTs, we tested four pig gut metagenomes (ERR1135178-ERR1135181) from [57]. According to myRT results, even higher proportions of DGR-RTs (41%-49%) were observed in these pig gut metagenomes (See Supplementary Table S4 and Supplementary Figure S1).

### Genomic context preferences of different classes of RTs

With predictions of putative RTs in complete genomes, we were able to identify domains that are frequently found in the proteins encoded by the neighboring genes of putative RTs (including those that are fused with the RT genes). Table 6 lists some of the co-occurring proteins/domains found in complete genomes (Supplementary Table S3 shows the frequent domains observed in the genomic neighborhood of RTs used in our RT collection, i.e., training data). Among non-redundant (identity *<* 90%) putative RTs in complete genomes, 86% of CRISPR-Cas RTs are found to co-occur with Cas1 domain, and 67% co-occur with Cas2. 61% of DGR-RTs are found to co-occur with Avd like domain; Avd like is found in bacterial accessory variability determinant (bAvd) proteins) in DGR systems. 77% of UG10 RTs are found together with SLATT 5 domain (families of SLATT domains are predicted to be associated with cell-suicide and diversity generating [58]). About 96% UG3 RTs are adjacent to a UG8 RT, and 93% of UG8-RTs are adjacent to a UG3 RT.

**Table 6:** Frequent domains encoded by the genes that are in the neighborhood of or fused to the genes encoding RTs in complete genomes.

| RT class | Domain | Co-occurrence frequency |
| --- | --- | --- |
| AbiA | HEPN_AbiA_CTD (appended) | 53% |
| CRISPR | Cas1 (Cas1_I-II-III,cas1,Cas_Cas1,cas1_HMARI,cas1_CYANO) | 86% |
| CRISPR | Cas2 | 69% |
| DGRs | Avd like | 61% |
| GII | GIIM (appended) | 58% |
| UG3 | UG8 | 96% |
| UG4 | Fimbrial domain (FimA, PRK15287, FimD, FimC, PRK15288) | 54% |
| UG5 | nitrilase | 39% |
| UG6 | group_II_RT_mat (appended) | 61% |
| UG8 | UG3 | 90% |
| UG9 | PRK14975 | 94% |
| UG10 | SLATT_5 | 76% |
| UG12 | AE_Prim_S_like/COG4951 | 71% |

Other domains encoded by the genes that are occasionally found in the neighborhood of RT genes include RelB (PF04221; antitoxin), RelB dinJ (antitoxin), dinJ-yafQ (toxin-antitoxin module), HTH XRE (Helix-turn-helix XRE-family), HTH_Tnp_1, Trypsin_2, AbiEii (Nucleotidyl transferase AbiEii toxin, Type IV TA system), DDE_Tnp_1 (transposase), metallo-hydrolase-like MBL-fold, mazF, xerC, AcrR (DNA-binding transcriptional regulator), AAA (ATPase family), SMC_prok_B, dnaG, DNA_pol_A, Phage integrase, InsE (Transposase and inactivated derivatives), T_den_put_tspse (putative transposase), and RAYT (REP element-mobilizing transposase), etc.

We observed nine reference genomes that have plasmids encoding group II introns RTs, and their adjacent genes are *bla*_*IMP* 26_ multi-drug resistance genes, which encode proteins containing_IMP_DIM-like_MBL-B1 domain (cd16301). We expanded our analysis and compiled a list of 25 plasmids that carry IMP_resistance genes and have a group II intron RT (see Supplementary Table S2). All of these group II intron RTs, have IMP DIM-like MBL-B1 in their genomic neighborhood except one (KX711880.1), and some also have Multi_Drug_Res (pfam PF00893) domain in their flanking genes. It seems that all of these group II intron RTs are almost identical to Kl.pn.I3 (ACJ76645.1). This result suggests the association of intron RTs and the mult-drug resistance (See Suplementary Table 2).

## DISCUSSIONS

In this study, we provided a tool for prediction, and classification of reverse transcriptase (RT) in bacterial genomes. Reverse transcriptases, the enzymes that convert RNA into cDNA, play substantial roles in different systems such as Diversity Generating Retroelements (DGRs), group II introns, CRISPR-Cas systems, retrons, etc. Identification of these RTs can provide us with information about the underlying interactions between phage and bacteria, archaea and archaeal viruses, and so forth. It can also help us to determine the origin of the RT, does it come from another species, or is it native (for instance DGR RTs that come from phage). Classification of RTs can also be extremely helpful when it comes to biotechnological/medical applications of certain classes of RTs, such as utilizing group II introns RTs as targetron and thermotargetron, and make use of retrons RTs in CRISPEY and SCRIBE methods, or as anti-phage defense systems. As RTs from each class have similar functions, characterization of every single RT is of importance, as it can shed light into identification of other RTs from the same class/family. Experimental studies can come to rescue, and identify the function of less-known/unknown groups of RTs. Thus, we provide a list of RTs in every single class (known/unknown), and even unclassified RTs in complete and bacterial genomes, as we believe these lists can be used by experimental and non-experimental researchers. (See Supplementary Table S1)

We expect that genomic neighborhood information can help provide insights into the putative function of unknown classes of RTs, including UG1-UG12, UN1-UN8, and CRISPR-like RTs. Fused domains in these RTs, alongside the information about the domains in the flanking genes of RTs in each class, can provide us with some insights into the functions of these RTs. Also, as we collect more data for each class, this information can be used or examined by experimental researchers. For example, SLATT 5 is frequently seen next to UG10 RTs. Our analyses show that 77% of RTs in complete genomes that have UG10 RT, also have SLATT 5 in the genomic neighborhood of the UG10 RT. Similarly, 67% of UG10 RTs in draft genomes have a SLATT 5 domain in their flanking genes. An example of SLATT domain next to a reverse transcriptase in *Salmonella enterica* subsp. enterica serovar 9,12:l,v:-str. 94293 is mentioned in [58]. This reverse transcriptase shares 92% identity with WP 015462025.1, UG10 RTs from *Edwardsiella piscicida* C07-087, which is mistakenly labeled as CRISPR RT in several other articles, yet our phylogenetic analysis showed that this RT groups with UG10 RTs, it has the SLATT 5 domain in its adjacent neighboring gene, and importantly we couldn’t detect any Cas neighbors, or CRISPR systems in this reference genome.

## DATA AVAILABILITY

MyRT is available to be used stand-alone (https://github.com/mgtools/myRT), and online (https://omics.sice.indiana.edu/myRT/). Predictions of RTs in reference genomes and selected metagenomes are available at https://omics.sice.indiana.edu/myRT/collection.php.

## References

[1] David Baltimore. Viral rna-dependent dna polymerase: Rna-dependent dna polymerase in virions of rna tumour viruses. Nature, 226(5252):1209–1211, 1970. [PubMed:4316300] [doi:10.1038/2261209a0].

[2] Stefan G Sarafianos, Bruno Marchand, Kalyan Das, Daniel M Himmel, Michael A Parniak, Stephen H Hughes, and Eddy Arnold. Structure and function of hiv-1 reverse transcriptase: molecular mechanisms of polymerization and inhibition. Journal of molecular biology, 385(3):693–713, 2009. [PubMed:19022262] [PubMed Central:PMC2881421] [doi:10.1016/j.jmb.2008.10.071].

[3] Thomas H Eickbush and Varuni K Jamburuthugoda. The diversity of retrotransposons and the properties of their reverse transcriptases. Virus research, 134(1-2):221–234, 2008. [PubMed:18261821] [PubMed Central:PMC2695964] [doi:10.1016/j.virusres.2007.12.010].

[4] Nicolás Toro and Rafael Nisa-Martínez. Comprehensive phylogenetic analysis of bacterial reverse transcriptases. PLoS One, 9(11):e114083, 2014. [PubMed:25423096] [PubMed Central:PMC4244168] [doi:10.1371/journal.pone.0114083].

[5] Fatemeh Sharifi and Yuzhen Ye. MyDGR: a server for identification and characterization of diversity-generating retroelements. Nucleic Acids Research, 47(W1):W289–W294, 2019. [PubMed:31049585] [PubMed Central:PMC6602519] [doi:10.1093/nar/gkz329].

[6] Steven Zimmerly and Li Wu. An unexplored diversity of reverse transcriptases in bacteria. Mobile DNA III, pages 1253–1269, 2015. [PubMed:26104699] [doi:10.1128/microbiolspec.MDNA3-0058-2014].

[7] Steven Zimmerly and Cameron Semper. Evolution of group ii introns. Mobile DNA, 6(1):7, 2015. [PubMed:25960782] [PubMed Central:PMC4424553] [doi:10.1186/s13100-015-0037-5].

[8] Peter J Enyeart, Georg Mohr, Andrew D Ellington, and Alan M Lambowitz. Biotechnological applications of mobile group ii introns and their reverse transcriptases: gene targeting, rna-seq, and non-coding rna analysis. Mobile DNA, 5(1):2, 2014. [PubMed:24410776] [PubMed Central:PMC3898094] [doi:10.1186/1759-8753-5-2].

[9] Fernando M García-Rodríguez, Teresa Hernández-Gutiérrez, Vanessa Díaz-Prado, and Nicolás Toro. Use of the computer-retargeted group ii intron rmint1 of sinorhizobium meliloti for gene targeting. RNA biology, 11(4):391–401, 2014. [PubMed:24646865] [PubMed Central:PMC4075523] [doi:10.4161/rna.28373].

[10] Georg Mohr, Wei Hong, Jie Zhang, Gu-zhen Cui, Yunfeng Yang, Qiu Cui, Ya-jun Liu, and Alan M Lambowitz. A targetron system for gene targeting in thermophiles and its application in clostridium thermocellum. PloS one, 8(7), 2013. [PubMed:23874856] [PubMed Central:PMC3706431] [doi:10.1371/journal.pone.0069032].

[11] Adi Millman, Aude Bernheim, Avigail Stokar-Avihail, Taya Fedorenko, Maya Voichek, Azita Leavitt, and Rotem Sorek. Bacterial retrons function in anti-phage defense. Cell, 2020. [doi:10.1101/2020.06.21.156273].

[12] Karen L Maxwell. Retrons: Complementing crispr in phage defense. The CRISPR Journal, 3(4):226–227, 2020. [PubMed:32833529] [doi:10.1089/crispr.2020.29100.kma].

[13] Jacob Bobonis, André Mateus, Birgit Pfalz, Sarela Garcia-Santamarina, Marco Galardini, Callie Kobayashi, Frank Stein, Mikhail M Savitski, Johanna R Elfenbein, Helene Andrews-Poymenis, et al. Bacterial retrons encode tripartite toxin/antitoxin systems. BioRxiv, 2020. [doi:10.1101/2020.06.22.160168].

[14] Jacob Bobonis, Karin Mitosch, André Mateus, George Kritikos, Johanna R Elfenbein, Mikhail M Savitski, Helene Andrews-Polymenis, and Athanasios Typas. Phage proteins block and trigger retron toxin/antitoxin systems. BioRxiv, 2020. [doi:10.1101/2020.06.22.160242].

[15] Anna J Simon, Andrew D Ellington, and Ilya J Finkelstein. Retrons and their applications in genome engineering. Nucleic acids research, 47(21):11007–11019, 2019. [PubMed:31598685] [PubMed Central:PMC6868368] [doi:10.1093/nar/gkz865].

[16] Sumit Handa, Yong Jiang, Sijia Tao, Robert Foreman, Raymond F Schinazi, Jeff F Miller, and Partho Ghosh. Template-assisted synthesis of adenine-mutagenized cdna by a retroelement protein complex. Nucleic acids research, 46(18):9711–9725, 2018. [PubMed:30007279] [PubMed Central:PMC6182149] [doi:10.1093/nar/gky620].

[17] Blair G Paul, Sarah C Bagby, Elizabeth Czornyj, Diego Arambula, Sumit Handa, Alexander Sczyrba, Partho Ghosh, Jeff F Miller, and David L Valentine. Targeted diversity generation by intraterrestrial archaea and archaeal viruses. Nature communications, 6(1):1–8, 2015. [PubMed:25798780] [PubMed Central:PMC4372165] [doi:10.1038/ncomms7585].

[18] Sergei Doulatov, Asher Hodes, Lixin Dai, Neeraj Mandhana, Minghsun Liu, Rajendar Deora, Robert W Simons, Steven Zimmerly, and Jeff F Miller. Tropism switching in bordetella bacteriophage defines a family of diversity-generating retroelements. Nature, 431(7007):476–481, 2004. [PubMed:15386016] [doi:10.1038/nature02833].

[19] Diego Arambula, Wenge Wong, Bob A Medhekar, Huatao Guo, Mari Gingery, Elizabeth Czornyj, Minghsun Liu, Sanghamitra Dey, Partho Ghosh, and Jeff F Miller. Surface display of a massively variable lipoprotein by a legionella diversity-generating retroelement. Proceedings of the National Academy of Sciences, 110(20):8212–8217, 2013. [PubMed:23633572] [PubMed Central:PMC3657778] [doi:10.1073/pnas.1301366110].

[20] Alec Vallota-Eastman, Eleanor C Arrington, Siobhan Meeken, Simon Roux, Krishna Dasari, Sydney Rosen, Jeff F Miller, David L Valentine, and Blair G Paul. Role of diversity-generating retroelements for regulatory pathway tuning in cyanobacteria. BMC genomics, 21(1):1–13, 2020. [PubMed:32977771] [PubMed Central:PMC7517822] [doi:10.1186/s12864-020-07052-5].

[21] Sean Benler, Ana Georgina Cobián-Güemes, Katelyn McNair, Shr-Hau Hung, Kyle Levi, Rob Edwards, and Forest Rohwer. A diversity-generating retroelement encoded by a globally ubiquitous bacteroides phage. Microbiome, 6(1):191, 2018. [PubMed:30352623] [PubMed Central:PMC6199706] [doi:10.1186/s40168-018-0573-6].

[22] Jeffrey K Cornuault, Marie-Agnès Petit, Mahendra Mariadassou, Leandro Benevides, Elisabeth Moncaut, Philippe Langella, Harry Sokol, and Marianne De Paepe. Phages infecting faecalibacterium prausnitzii belong to novel viral genera that help to decipher intestinal viromes. Microbiome, 6(1):65, 2018. [PubMed:29615108] [PubMed Central:PMC5883640] [doi:10.1186/s40168-018-0452-1].

[23] Yuzhen Ye. Identification of diversity-generating retroelements in human microbiomes. International journal of molecular sciences, 15(8):14234–14246, 2014. [PubMed:25196521] [PubMed Central:PMC4159848] [doi:10.3390/ijms150814234].

[24] Nicolas Toro, Mario Rodríguez Mestre, Francisco Martínez-Abarca, and Alejandro González-Delgado. Recruitment of reverse transcriptase-cas1 fusion proteins by type vi-a crispr-cas systems. Frontiers in microbiology, 10:2160, 2019. [PubMed:31572350] [doi:10.3389/fmicb.2019.02160].

[25] Nicolás Toro, Francisco Martínez-Abarca, and Alejandro González-Delgado. The reverse transcriptases associated with crispr-cas systems. Scientific reports, 7(1):1–7, 2017. [PubMed:28769116] [PubMed Central:PMC5541045] [doi:10.1038/s41598-017-07828-y].

[26] Nicolás Toro, Francisco Martínez-Abarca, Mario Rodríguez Mestre, and Alejandro González-Delgado. Multiple origins of reverse transcriptases linked to crispr-cas systems. RNA biology, 16(10):1486–1493, 2019. [PubMed:31276437] [PubMed Central:PMC6779382] [doi:10.1080/15476286.2019.1639310].

[27] Sukrit Silas, Kira S Makarova, Sergey Shmakov, David Páez-Espino, Georg Mohr, Yi Liu, Michelle Davison, Simon Roux, Siddharth R Krishnamurthy, Becky Xu Hua Fu, et al. On the origin of reverse transcriptase-using crispr-cas systems and their hyperdiverse, enigmatic spacer repertoires. MBio, 8(4), 2017. [PubMed:28698278] [PubMed Central:PMC5513706] [doi:10.1128/mBio.00897-17].

[28] Louis-Charles Fortier, Julie D Bouchard, and Sylvain Moineau. Expression and sitedirected mutagenesis of the lactococcal abortive phage infection protein abik. Journal of bacteriology, 187(11):3721–3730, 2005. [PubMed:15901696] [PubMed Central:PMC1112063] [doi:10.1128/JB.187.11.3721-3730.2005].

[29] Richard Odegrip, Anders S Nilsson, and Elisabeth Haggård-Ljungquist. Identification of a gene encoding a functional reverse transcriptase within a highly variable locus in the p2-like coliphages. Journal of bacteriology, 188(4):1643–1647, 2006. [PubMed:16452449] [PubMed Central:PMC1367236] [doi:10.1128/JB.188.4.1643-1647.2006].

[30] Kimberley D Seed. Battling phages: how bacteria defend against viral attack. PLoS pathogens, 11(6), 2015. [PubMed:26066799] [PubMed Central:PMC4465916] [doi:10.1371/journal.ppat.1004847].

[31] Vivek Anantharaman, Kira S Makarova, A Maxwell Burroughs, Eugene V Koonin, and L Aravind. Comprehensive analysis of the hepn superfamily: identification of novel roles in intra-genomic conflicts, defense, pathogenesis and rna processing. Biology direct, 8(1):15, 2013. [PubMed:23768067] [PubMed Central:PMC3710099] [doi:10.1186/1745-6150-8-15].

[32] Marie-Christine Chopin, Alain Chopin, and Elena Bidnenko. Phage abortive infection in lactococci: variations on a theme. Current opinion in microbiology, 8(4):473–479, 2005. [PubMed:15979388] [doi:10.1016/j.mib.2005.06.006].

[33] Dawn M Simon and Steven Zimmerly. A diversity of uncharacterized reverse transcriptases in bacteria. Nucleic acids research, 36(22):7219–7229, 2008. [PubMed:19004871] [PubMed Central:PMC2602772] [doi:10.1093/nar/gkn867].

[34] Linyi Gao, Han Altae-Tran, Francisca Böhning, Kira S Makarova, Michael Segel, Jonathan L Schmid-Burgk, Jeremy Koob, Yuri I Wolf, Eugene V Koonin, and Feng Zhang. Diverse enzymatic activities mediate antiviral immunity in prokaryotes. Science, 369(6507):1077–1084, 2020. [PubMed:32855333] [doi:10.1126/science.aba0372].

[35] Manuel A Candales, Adrian Duong, Keyar S Hood, Tony Li, Ryan AE Neufeld, Runda Sun, Bonnie A McNeil, Li Wu, Ashley M Jarding, and Steven Zimmerly. Database for bacterial group ii introns. Nucleic acids research, 40(D1):D187–D190, 2012. [PubMed:22080509] [PubMed Central:PMC3245105] [doi:10.1093/nar/gkr1043].

[36] Michael Abebe, Manuel A Candales, Adrian Duong, Keyar S Hood, Tony Li, Ryan AE Neufeld, Abat Shakenov, Runda Sun, Li Wu, Ashley M Jarding, et al. A pipeline of programs for collecting and analyzing group ii intron retroelement sequences from genbank. Mobile DNA, 4(1):28, 2013. [doi:10.1186/1759-8753-4-28].

[37] Li Wu, Mari Gingery, Michael Abebe, Diego Arambula, Elizabeth Czornyj, Sumit Handa, Hamza Khan, Minghsun Liu, Mechthild Pohlschroder, Kharissa L Shaw, et al. Diversity-generating retroelements: natural variation, classification and evolution inferred from a large-scale genomic survey. Nucleic acids research, 46(1):11–24, 2018. [PubMed:29186518] [PubMed Central:PMC5758913] [doi:10.1093/nar/gkx1150].

[38] Thomas Schillinger and Nora Zingler. The low incidence of diversity-generating retroelements in sequenced genomes. Mobile genetic elements, 2(6):287–291, 2012. [PubMed:23481467] [PubMed Central:PMC3575424] [doi:10.4161/mge.23244].

[39] Samuel Minot, Alexandra Bryson, Christel Chehoud, Gary D Wu, James D Lewis, and Frederic D Bushman. Rapid evolution of the human gut virome. Proceedings of the National Academy of Sciences, 110(30):12450–12455, 2013. [PubMed:23836644] [PubMed Central:PMC3725073] [doi:10.1073/pnas.1300833110].

[40] Curtis Huttenhower, Dirk Gevers, Rob Knight, Sahar Abubucker, Jonathan H Badger, Asif T Chinwalla, Heather H Creasy, Ashlee M Earl, Michael G FitzGerald, Robert S Fulton, et al. Structure, function and diversity of the healthy human microbiome. nature, 486(7402):207, 2012. [PubMed:22699609] [PubMed Central:PMC3564958] [doi:10.1038/nature11234].

[41] Shennan Lu, Jiyao Wang, Farideh Chitsaz, Myra K Derbyshire, Renata C Geer, Noreen R Gonzales, Marc Gwadz, David I Hurwitz, Gabriele H Marchler, James S Song, et al. Cdd/sparcle: the conserved domain database in 2020. Nucleic acids research, 48(D1):D265–D268, 2020. [PubMed:31777944] [PubMed Central:PMC6943070] [doi:10.1093/nar/gkz991].

[42] Weizhong Li and Adam Godzik. Cd-hit: a fast program for clustering and comparing large sets of protein or nucleotide sequences. Bioinformatics, 22(13):1658–1659, 2006. [PubMed:16731699] [doi:10.1093/bioinformatics/btl158].

[43] Sean R Eddy. Accelerated profile hmm searches. PLoS Comput Biol, 7(10):e1002195, 2011. [PubMed:22039361] [PubMed Central:PMC3197634] [doi:10.1371/journal.pcbi.1002195].

[44] Sara El-Gebali, Jaina Mistry, Alex Bateman, Sean R Eddy, Aurélien Luciani, Simon C Potter, Matloob Qureshi, Lorna J Richardson, Gustavo A Salazar, Alfredo Smart, et al. The pfam protein families database in 2019. Nucleic acids research, 47(D1):D427–D432, 2019. [PubMed:30357350] [PubMed Central:PMC6324024] [doi:10.1093/nar/gky995].

[45] Robert C Edgar. Muscle: a multiple sequence alignment method with reduced time and space complexity. BMC bioinformatics, 5(1):113, 2004. [PubMed:15318951] [PubMed Central:PMC517706] [doi:10.1186/1471-2105-5-113].

[46] Morgan N Price, Paramvir S Dehal, and Adam P Arkin. Fasttree 2–approximately maximum-likelihood trees for large alignments. PloS one, 5(3):e9490, 2010. [PubMed:20224823] [PubMed Central:PMC2835736] [doi:10.1371/journal.pone.0009490].

[47] Sean R. Eddy. Profile hidden markov models. Bioinformatics (Oxford, England), 14(9):755–763, 1998. [PubMed:9918945] [doi:10.1093/bioinformatics/14.9.755].

[48] Mina Rho, Haixu Tang, and Yuzhen Ye. Fraggenescan: predicting genes in short and error-prone reads. Nucleic acids research, 38(20):e191–e191, 2010. [PubMed:20805240] [PubMed Central:PMC2978382] [doi:10.1093/nar/gkq747].

[49] Frederick A Matsen, Robin B Kodner, and E Virginia Armbrust. pplacer: linear time maximum-likelihood and bayesian phylogenetic placement of sequences onto a fixed reference tree. BMC bioinformatics, 11(1):538, 2010. [PubMed:21034504] [PubMed Central:PMC3098090] [doi:10.1186/1471-2105-11-538].

[50] Li-Gen Wang, Tommy Tsan-Yuk Lam, Shuangbin Xu, Zehan Dai, Lang Zhou, Tingze Feng, Pingfan Guo, Casey W Dunn, Bradley R Jones, Tyler Bradley, et al. treeio: an r package for phylogenetic tree input and output with richly annotated and associated data. Molecular biology and evolution, 37(2):599–603, 2020. [PubMed:31633786] [PubMed Central:PMC6993851] [doi:10.1093/molbev/msz240].

[51] Stilianos Louca and Michael Doebeli. Efficient comparative phylogenetics on large trees. Bioinformatics, 34(6):1053–1055, 2018. [PubMed:29091997] [doi:10.1093/bioinformatics/btx701].

[52] Anthony M Bolger, Marc Lohse, and Bjoern Usadel. Trimmomatic: a flexible trimmer for illumina sequence data. Bioinformatics, 30(15):2114–2120, 2014. [PubMed:24695404] [PubMed Central:PMC4103590] [doi:10.1093/bioinformatics/btu170].

[53] Quan Zhang and Yuzhen Ye. Not all predicted crispr–cas systems are equal: isolated cas genes and classes of crispr like elements. BMC bioinformatics, 18(1):92, 2017. [PubMed:28166719] [PubMed Central:PMC5294841] [doi:10.1186/s12859-017-1512-4].

[54] Mira V Han and Christian M Zmasek. phyloxml: Xml for evolutionary biology and comparative genomics. BMC bioinformatics, 10(1):356, 2009. [PubMed:19860910] [PubMed Central:PMC2774328] [doi:10.1186/1471-2105-10-356].

[55] Kenji K Kojima and Minoru Kanehisa. Systematic survey for novel types of prokaryotic retroelements based on gene neighborhood and protein architecture. Molecular biology and evolution, 25(7):1395–1404, 2008. [PubMed:18391066] [doi:10.1093/molbev/msn081].

[56] Sunghee Lee, Brandi Cantarel, Bernard Henrissat, Dirk Gevers, Bruce W Birren, Curtis Huttenhower, and GwangPyo Ko. Gene-targeted metagenomic analysis of glucan-branching enzyme gene profiles among human and animal fecal microbiota. The ISME journal, 8(3):493–503, 2014. [PubMed:24108330] [PubMed Central:PMC3930310] [doi:10.1038/ismej.2013.167].

[57] Liang Xiao, Jordi Estellé, Pia Kiilerich, Yuliaxis Ramayo-Caldas, Zhongkui Xia, Qiang Feng, Suisha Liang, Anni Øyan Pedersen, Niels JØrgen Kjeldsen, Chuan Liu, et al. A reference gene catalogue of the pig gut microbiome. Nature microbiology, 1(12):1–6, 2016. [PubMed:27643971] [doi:10.1038/nmicrobiol.2016.161].

[58] A Maxwell Burroughs, Dapeng Zhang, Daniel E Schäffer, Lakshminarayan M Iyer, and L Aravind. Comparative genomic analyses reveal a vast, novel network of nucleotide-centric systems in biological conflicts, immunity and signaling. Nucleic acids research, 43(22):10633–10654, 2015. [PubMed:26590262] [PubMed Central:PMC4678834] [doi:10.1093/nar/gkv1267].

