## Supplementary Tables and Figures for "Identification and Classification of Reverse Transcriptases in Bacterial Genomes and Metagenomes"

### Supplementary materials: Identification and Characterization of Reverse Transcriptases in Bacterial Genomes and Metagenomes

Fatemeh Sharifi[1] and Yuzhen Ye[2,*]

[1,2]*Luddy School of Informatics, Computing, and Engineering , Indiana University, Bloomington, IN 47408, USA*

Table 1: Distribution of different classes of RT in complete and draft genomes*

| Index | RT Class | In Complete Genomes | In Draft Genomes | Sum |
|---|---|---|---|---|
| 1 | AbiA | 33 | 294 | 327 |
| 2 | AbiK | 102 | 2914 | 3016 |
| 3 | AbiP2 | 227 | 2836 | 3063 |
| 4 | CRISPR (CRISPR RT w/ Cas genes) | 140 | 2026 | 2166 |
| 5 | CRISPR-like | 199 | 3962 | 4161 |
| 6 | CRISPR-like2 (CRISPR RT w/o Cas genes) | 125 | 1973 | 2098 |
| 7 | CRISPR-like3 (Other RT w/ Cas genes) | 67 | 19 | 86 |
| 8 | DGRs | 459 | 16320 | 16779 |
| 9 | G2L4 | 29 | 514 | 543 |
| 10 | GII (Group II Introns) | 18761 | 163257 | 182018 |
| 11 | Retrons | 3102 | 34259 | 37361 |
| 12 | UG1 | 71 | 685 | 756 |
| 13 | UG2 | 398 | 3104 | 3502 |
| 14 | UG3 | 196 | 1589 | 1785 |
| 15 | UG4 | 226 | 2374 | 2600 |
| 16 | UG5 | 212 | 2690 | 2902 |
| 17 | UG6 | 109 | 101 | 210 |
| 18 | UG7 | 129 | 1152 | 1281 |
| 19 | UG8 | 213 | 2317 | 2530 |
| 20 | UG9 | 29 | 295 | 324 |
| 21 | UG10 | 178 | 2015 | 2193 |
| 22 | UG12 | 23 | 214 | 237 |
| 23 | UN1 | 43 | 469 | 512 |
| 24 | UN2 | 17 | 214 | 231 |
| 25 | UN3 | 88 | 719 | 807 |
| 26 | UN4 | 85 | 1157 | 1242 |
| 27 | UN5 | 26 | 186 | 212 |
| 28 | UN6 | 47 | 440 | 487 |
| 29 | UN7 | 22 | 253 | 275 |
| 30 | UN8 | 23 | 173 | 196 |
| 31 | UNC | 172 | 3361 | 3533 |
| 32 | Sum | 25484 | 251863 | 277347 |

* As of 10/22/2020

Table 2: Plasmids harbouring IMP-4 and IMP-26* [1, 2]

| Accession NO. | Plasmid | Carbapenemase | Source | Length(bp) | myRT results |
|---|---|---|---|---|---|
| KM977631.1 | pIMP-1495 | IMP-4 | *Klebsiella Pneumoniae* | 50742 | myRT prediction |
| KT982615.1 | pIMP-FS1505 | IMP-4 | *Escherichia coli* | 54449 | myRT prediction |
| KT989598.1 | pIMP-SH1506 | IMP-4 | *Enterobacter cloacae* | 54669 | myRT prediction |
| KU051708.1 | pIMP-SZ1501 | IMP-4 | *Klebsiella Pneumoniae* | 51469 | myRT prediction |
| KU051710.1 | pIMP-FJ1503 | IMP-4 | *Citrobacter freundii* | 50546 | myRT prediction |
| CP028486.1 | p3 | IMP-4 | *Escherichia coli* | 52864 | myRT prediction |
| KU862632.1 | pIMP-KP1495 | IMP-4 | *Klebsiella Pneumoniae* | 51591 | myRT prediction |
| KX711879.1 | P378-IMP | IMP-4 | *Pseudomonas aeruginosa* | 51207 | myRT prediction |
| KY913900.1 | p4-IPM | IMP-4 | *Klebsiella oxytoca* | 61680 | myRT prediction |
| MF344559.1 | p128379-IMP | IMP-4 | *Enterobacter hormaechei* | 42279 | myRT prediction |
| CP033103.1 | pEHZJ1 | IMP-26 | *Enterobacter hormaechei* | 343918 | myRT prediction |
| MH399264.1 | pIMP26 | IMP-26 | *Enterobacter cloacae* | 329420 | myRT prediction |

More examples are available at myRT-blaIMP

Table 3: Frequent domains in the genomic neighborhood of previously labeled RTs

| RT class | Frequent neighbors |
|---|---|
| CRISPR-Cas RTs | Cas1 and Cas2 |
| DGRs RTs | Avd_like, YfmG, and DUF1566 |
| GII RTs | GIMM and RVT_N |
| UG10 RTs | SLATT_5 (88%) |
| UG3 RTs | UG8 RTs (100%) |
| UG8 RTs | UG3 RTs (91%) |
| UG9 RTs | PRK14975 * (75%) |
| UG12 RTs | AE_Prim_S_like (appended; 100%) |

*PRK14975: bifunctional 3'-5' exonuclease/DNA polymerase

Table 4: myRT results for different metagenomes

| Metagenome | Source | myRT results | myDGR results |
|---|---|---|---|
| ERR248260 | Chicken | myRT prediction | myDGR prediction |
| ERR248261 | Cow | myRT prediction | myDGR prediction |
| ERR248262 | Human | myRT prediction | myDGR prediction |
| ERR248263 | Pig | myRT prediction | myDGR prediction |
| ERR1135178 | Pig | myRT prediction | myDGR prediction |
| ERR1135179 | Pig | myRT prediction | myDGR prediction |
| ERR1135180 | Pig | myRT prediction | myDGR prediction |
| ERR1135181 | Pig | myRT prediction | myDGR prediction |

Table 5: myRT results for individual genomes

| Accession | Genome/Plasmid | myRT results |
|---|---|---|
| CP005935.1 | *Bacillus thuringiensis* YBT-1518 | myRT prediction |
| BA000019.2 | *Nostoc sp.* PCC 7120 | myRT prediction |
| AP009552.1 | *Microcystis aeruginosa* NIES-843 | myRT prediction |
| U17233.3 | *Lactococcus lactis* plasmid pTR2030 | myRT prediction |
| U35629.2 | *Lactococcus lactis* plasmid pSRQ800 | myRT prediction |
| CP000247.1 | *Escherichia coli* 536 | myRT prediction |
| JGYE01000047.1 | *Salmonella enterica* subsp. enterica serovar 9,12:l,v:- str. 94293 | myRT prediction |
| CP004141.1 | Edwardsiella piscicida C07-087 | myRT prediction |
| FP929062.1 | *Clostridiales sp.* SS3/4 (draft genome) | myRT prediction |

Table 6: Potential improvements of RT classification by using phylogenetic information

| Accession number | Gene coordinates | hmmscan | pplacer | genomic neighborhood | myRT results |
|---|---|---|---|---|---|
| ACK61740.1 | CP001176.1_543382_544635_+ | **UG3**/Retrons/DGRs | UG3 | UG8 | myRT prediction |
| ADE85032.1 | CP001312.1_1357974_1358504_- | GII/**CRISPR** | CRISPR | CRISPR_Cas6 | myRT prediction |
| - | KB901875.1_2019009_2019602_+ | CRISPR/**GII** | GII | GIIM | myRT prediction |
| ANU66363.2 | CP015403.2_1765974_1766285_- | DGRs/**GII** | GII | GIIM | myRT prediction |
| ABW11582.1 | CP000820.1_2576306_2576710_+ | **GII**/CRISPR | GII | GIIM | myRT prediction |
| ACN15726.1 | CP001087.1_3000453_3001124_- | DGRs/**GII** | GII | GII | myRT prediction |
| BAZ36932.1 | AP018280.1_22270_22950_+ | DGRs/**GII** | GII | McrA | myRT prediction |
| ABW09889.1 | CP000820.1_498987_499841_+ | DGRs/**GII** | GII | INT_RitC_C_like | myRT prediction |
| BAQ13887.1 | AP014696.1_2236430_2239687_- | UG10/Retrons/**UG6** | UG6 | HNH_4 | myRT prediction |
| ACA53940.1 | CP000962.1_2352044_2355301_- | UG10/Retrons/**UG6** | UG6 | HNH_4 | myRT prediction |
| BBI33370.1 | AP019400.1_3194415_3197627_+ | DGRs/**UG6**/Retrons | UG6 | nitrilase | myRT prediction |
| AIG26831.1 | CP007806.1_2784436_2786124_- | UN4/**AbiK**/AbiP2 | AbiK | HTH_21, InsE | myRT prediction |
| QCQ34047.1 | CP037440.1_5056385_5059810_+ | UG6/**Retrons**/UG1 | Retrons | dnaG | myRT prediction |
| AXJ12401.1 | CP022601.1_438418_439032_+ | UN1/**UG7**/CRISPR | UG7 | UG7, InsE | myRT prediction |
| AXL52307.1 | CP031467.1_2031852_2033405_- | DGRs/Retrons/**CRISPR-like** | CRISPR-like | CytC5 | myRT prediction |

Table 7: Re-classification of 19 of the previously labeled RTs

| Accession number | Old classification | New classification | genomic neighborhood | myRT results |
|---|---|---|---|---|
| CBL40120.1 | UNC | DGRs | Phage_XkdX | myRT prediction |
| ZP_01872295.1 (EDM23124.1) | UG11 | CRISPR-Cas | *Cas1* | myRT prediction |
| EGP13976.1 (AEB93977.1) | DGRs | Group II introns | GIIM[c] | myRT prediction |
| YP_001397265.1 (EDK35894.1) | DGRs | Group II introns | Intron_maturas2[c] | myRT prediction |
| AFZ16538.1 (WP_015180701.1) | UNC | AbiA | MazF, HTH_XRE | myRT prediction |
| NP_442332.1 | UG3 | UG7 | | myRT prediction |
| AFY59940.1 | UG3 | UG7 | | myRT prediction |
| AGA07305.1 | UG6 | UN3 | | myRT prediction |
| EPZ72367.1 | UNC | UN4 | | myRT prediction |
| AGI67543.1 | Group II like 3[a] | UG3 | UG8 | myRT prediction |
| AEJ99900.1 | Group II like 4[a] | UG4 | FimD, FimA | myRT prediction |
| AGK68212.1 (YP_005196274.1) | UG11[a] | Retrons | | myRT prediction |
| CCF10237.1 (EQR96236) | UG11[a] | Retrons | | myRT prediction |
| WP_007781002.1 (EJL44959.1) | CRISPR-Cas[b] | DGRs | Avd_like | myRT prediction |
| WP_015462025.1 | CRISPR-Cas[b] | UG10 | SLATT_5 | myRT prediction |
| WP_036415250.1 | CRISPR-Cas[b] | UG10 | SLATT_5 | myRT prediction |

[*] Even though the reference genome has RT(s), these genes have no RT domain.
[a] These entries are from [3].[b] These labels are based on [4]. [c] These domains are fused to the gene on column 1.

Table 8: Result of testing myRT on the datasets from [3]

| Accession number | Gene coordinates | Class | myRT Prediction | myRT results |
|---|---|---|---|---|
| CAA78293.1 | AHAX01000006.1_19205_20143_+ | Retrons | Retrons | myRT prediction |
| AAM42896.1 | AE008922.1_4324395_4326089_+ | Retrons | Retrons | myRT prediction |
| AFY43443.1 | CP003548.1_3308170_3310188_- | CRISPR* | CRISPR | myRT prediction |
| ZP_01854760.1 | NZ_ABCE01000016.1_73333_76125_+ | G2L5 | GII | myRT prediction |
| ZP_01851752.1 | ABCE01000001.1_117632_117919_- | G2L5 | CRISPR-like2 | myRT prediction |
| EKP98429.1 | CP006690.1_1914333_1915748_- | UG2 | UG2 | myRT prediction |
| CCC73043.1 | HE576794.1_939599_941005_+ | UG2 | UG2 | myRT prediction |
| AEG09910.1 | CP002767.1_716105_717388_ | UG2 | UG2 | myRT prediction |
| AEW72297.1 | CP002886.1_903224_904396_+ | UG3 | UG3 | myRT prediction |
| ACD38707.1 | EU595736.1_18886_20160_+ | UG3 | UG3 | myRT prediction |
| AHE72406.1 | CP006580.1_34213_36348_- | UG4 | UG4 | myRT prediction |
| AGI74246.1 | CP003742.1_4621630_4622688_+ | UG4 | UG4 | myRT prediction |
| CDI94624.1 | HG530068.1_6882792_6885812_+ | UG5 | UG5 | myRT prediction |
| AHM47056.1 | CP007393.1_928229_931438_- | UG5 | UG5 | myRT prediction |
| ACU61669.1 | CP001699.1_5262939_5266187_+ | UG6 | UG6 | myRT prediction |
| AHL77683.1 | CP007441.1_3859031_3861010_- | UG8 | UG8 | myRT prediction |
| AEN62840.1 | CP003026.1_82890_84845_- | UG8 | UG8 | myRT prediction |
| AFG37103.1 | CP003282.1_1170484_1172610_- | UG8 | UG8 | myRT prediction |
| ADI30165.1 | CP002056.1_2016134_2017783_- | UG9 | UG9 | myRT prediction |
| AGA65410.1 | CP002873.1_44531_46162_+ | UN5# | UN5 | myRT prediction |

* We note that AFY43443.1 was labeled as a G2L1/G2L2 RT in [3], but the authors reported that the RTs classified as G2L1 and G2L2 are associated with *cas1* genes of CRISPR/Cas loci. G2L1 and G2L2 were proposed originally in [5], and in Toro et al [6], they were renamed as CRISPR-RT because of their association with the CRISPR-Cas systems.# AGA65410.1: this RT was classified as UG14 in [3], which includes three UG14 RTs including AGA65419.1, BAL31358.1 and AEA43024.1; all these three sequences were classified as UN5 by myRT, which is consistent with Toro et al's classification, which classified BAL31358.1 as a UN5 RT [6].

Table 9: Evaluation of myRT on the collection of CRISPR-Cas RTs from [7]

| Genome | RT coordinates# | myRT prediction | identity%& | myRT results |
|---|---|---|---|---|
| ASPN01000006.1 | ASPN01000006.1_20247_22901_+ | CRISPR | 34 | myRT prediction |
| AQRP01000065.1 | AQRP01000065.1_13318_16137_+ | CRISPR | 34 | myRT prediction |
| JXXW01000010.1 | JXXW01000010.1_71385_73319_+ | CRISPR | 59 | myRT prediction |
| ASAJ01000015.1 | ASAJ01000015.1_125843_128860_- | CRISPR | 56 | myRT prediction |
| KL370780.1 | KL370780.1_93557_95149_+ | CRISPR | 35 | myRT prediction |
| JWIO01000025.1 | JWIO01000025.1_10124_11221_+ | CRISPR | 38 | myRT prediction |
| BCQS01000026.1 | BCQS01000026.1_4694_5707_- | CRISPR | 40 | myRT prediction |
| JH470356.1 | JH470356.1_27166_28113_+ | CRISPR | 77 | myRT prediction |
| LLVU01000015.1 | LLVU01000015.1_5740_7752_- | CRISPR | 57 | myRT prediction |
| LOAS01000028.1 | LOAS01000028.1_37027_38064_- | CRISPR | 48 | myRT prediction |
| JYJP01000030.1 | JYJP01000030.1_21534_23426_+ | CRISPR | 74 | myRT prediction |
| JH992891.1 | JH992891.1_86001_86924_- | CRISPR | 75 | myRT prediction |
| ANNX01000115.1 | ANNX01000115.1_14418_15371_- | CRISPR | 76 | myRT prediction |
| ALWD01000163.1 | ALWD01000163.1_24045_25013_+ | CRISPR | 84.62 | myRT prediction |
| ANNX01000117.1 | ANNX01000117.1_21795_22772_- | CRISPR | 86 | myRT prediction |
| JH992901.1 | JH992901.1_1016250_1017311_+ | CRISPR | 81 | myRT prediction |
| JXCA01000005.1 | JXCA01000005.1_398870_399850_- | CRISPR | 82 | myRT prediction |
| ALVY01000183.1 | ALVY01000183.1_27387_28304_+ | CRISPR | 69 | myRT prediction |
| HE972669.1 | HE972669.1_45936_46913_- | CRISPR | 72 | myRT prediction |
| ALWB01000016.1 | ALWB01000016.1_12151_14244_- | CRISPR | 64 | myRT prediction |
| ASMA01000004.1 | ASMA01000004.1_21607_22458_+ | CRISPR | 46 | myRT prediction |
| JQFA01000004.1 | JQFA01000004.1_1149670_1151637_- | CRISPR | 85 | myRT prediction |
| AJLK01000155.1 | AJLK01000155.1_941_2959_- | CRISPR | 86 | myRT prediction |
| ASZN01000033.1 | ASZN01000033.1_1862_3346_- | CRISPR | 52 | myRT prediction |
| LAQJ01000220.1 | LAQJ01000220.1_7006_7917_- | CRISPR | 37 | myRT prediction |
| KK211136.1 | KK211136.1_10156_11052_- | CRISPR | 62 | myRT prediction |
| BAFN01000001.1 | BAFN01000001.1_294229_295173_+ | CRISPR-like2 | 71 | myRT prediction |
| AJKO01000007.1 | AJKO01000007.1_124518_125894_- | UG2* | 34 (UG2) | myRT prediction |
| DS570667.1 | DS570667.1_23830_25101_- | UNC* | 29 | myRT prediction |
| CP007699.1 | CP007699.1_7617452_7618696_+ | CRISPR-like* | 38 (GII) | myRT prediction |
| LAKD01000050.1 | LAKD01000050.1_46946_48151_- | CRISPR-like* | 25 | myRT prediction |

* These 4 RTs were referred as ”RTs with unusual RT-associated CRISPR-Cas architectures” in [7]. #: Coordinates of the predicted RT genes are presented in the genome/contig_start_end_strand format, which shows the start, end, and strand of the protein coding regions. & This column lists the highest sequence identity between the predicted RT and the RVT_1 domains used to build the HMMs for the different classes of RTs.

Table 10: Evaluation of myRT on the collection of retron RTs from [8], all of which were predicted as retron RT by myRT.

| Known Retron | Genome | RT coordinates | Identity%[&] | myRT results |
|---|---|---|---|---|
| EC48[a] | *Escherichia Coli* DE147 | LFQP01000005.1_154506_155696_- | 50 | myRT prediction |
| EC67[a] | *Escherichia coli* S10 | CP010229.1_4712073_4713833_- | 61 | myRT prediction |
| EC73[a] | *Escherichia coli* M10 | CP010200.1_2393178_2394128_+ | 36 | myRT prediction |
| Ec78[a] | *Escherichia coli* 102598 | JHRW01000018.1_27622_28557_+ | 49 | myRT prediction |
| EC83[a] | *Escherichia coli* 05-2753 | CXYK01000012.1_74586_75524_+ | 47 | myRT prediction |
| Mx65 | *Myxococcus xanthus* DSM 16526 | FNOH01000027.1_37959_39242_+ | 53 | myRT prediction |
| Eco8[a] | *Escherichia coli* 200499 | CYGJ01000003.1_369367_370491_+ | 47 | myRT prediction |
| Se72 | *Salmonella enterica*[b] | AMMS01000284.1_2640_3671_- | 49 | myRT prediction |
| Vc137[a] | *Vibrio cholerae* 2012EL-1759 | JNEW01000012.1_609188_610135_+ | 49 | myRT prediction |
| Vp96 | *Vibrio parahaemolyticus* S119 | AWJG01000250.1_32_1054_+ | 49 | myRT prediction |
| YF79 | *Yersinia frederiksenii* ATCC 33641 | KN150731.1_1692670_1693602_- | 50 | myRT prediction |

[a] These retrons function as anti-phage defense systems. [b] Salmonella enterica enterica sv. Heidelberg 579083-10. [&] This column lists the highest sequence identity between the predicted RT and the RVT_1 domains used to build the HMMs for the different classes of RTs.

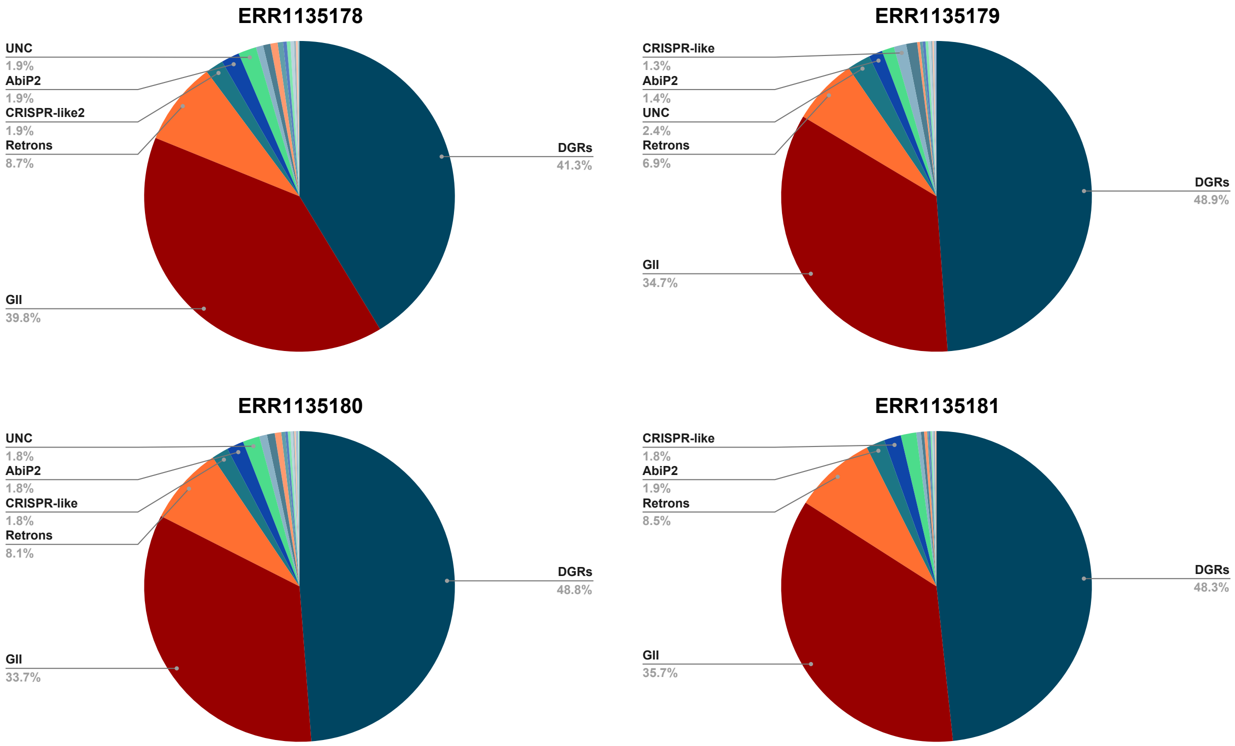


Figure 1: Distribution of different RT classes in pig gut metagenomes from [9]

### References

[1] Yingying Hao, Chunhong Shao, Xu Geng, Yuanyuan Bai, Yan Jin, and Zhiming Lu. Genotypic and phenotypic characterization of clinical escherichia coli sequence type 405 carrying incn2 plasmid harboring blandm-1. *Frontiers in microbiology*, 10:788, 2019. [PubMed:31105653] [PubMed Central:PMC6499153] [doi:10.3389/fmicb.2019.00788].

[2] Jian-Jun Gou, Na Liu, Li-Hua Guo, Hao Xu, Tao Lv, Xiao Yu, Yun-Bo Chen, Xiao-Bing Guo, Yu-Ting Rao, and Bei-Wen Zheng. Carbapenem-resistant enterobacter hormaechei st1103 with imp-26 carbapenemase and esbl gene blashv-178. *Infection and Drug Resistance*, 13:597, 2020. [PubMed:32110070] [PubMed Central:PMC7039083] [doi:10.2147/IDR.S232514].

[3] Steven Zimmerly and Li Wu. An unexplored diversity of reverse transcriptases in bacteria. *Mobile DNA III*, pages 1253–1269, 2015. [PubMed:26104699] [doi:10.1128/microbiolspec.MDNA3-0058-2014].

[4] Nicolás Toro, Francisco Martínez-Abarca, and Alejandro González-Delgado. The reverse transcriptases associated with crispr-cas systems. *Scientific reports*, 7(1):1–7, 2017. [PubMed:28769116] [PubMed Central:PMC5541045] [doi:10.1038/s41598-017-07828-y].

[5] Dawn M Simon and Steven Zimmerly. A diversity of uncharacterized reverse transcriptases in bacteria. *Nucleic acids research*, 36(22):7219–7229, 2008. [PubMed:19004871] [PubMed Central:PMC2602772] [doi:10.1093/nar/gkn867].

[6] Nicolás Toro and Rafael Nisa-Martínez. Comprehensive phylogenetic analysis of bacterial reverse transcriptases. *PLoS One*, 9(11):e114083, 2014. [PubMed:25423096] [PubMed Central:PMC4244168] [doi:10.1371/journal.pone.0114083].

[7] Sukrit Silas, Kira S Makarova, Sergey Shmakov, David Páez-Espino, Georg Mohr, Yi Liu, Michelle Davison, Simon Roux, Siddharth R Krishnamurthy, Becky Xu Hua Fu, et al. On the origin of reverse transcriptase-using crispr-cas systems and their hyperdiverse, enigmatic spacer repertoires. *MBio*, 8(4), 2017. [PubMed:28698278] [PubMed Central:PMC5513706] [doi:10.1128/mBio.00897-17].

[8] Anna J Simon, Andrew D Ellington, and Ilya J Finkelstein. Retrons and their applications in genome engineering. *Nucleic acids research*, 47(21):11007–11019, 2019. [PubMed:31598685] [PubMed Central:PMC6868368] [doi:10.1093/nar/gkz865].

[9] Liang Xiao, Jordi Estellé, Pia Kiilerich, Yuliaxis Ramayo-Caldas, Zhongkui Xia, Qiang Feng, Suisha Liang, Anni Øyan Pedersen, Niels Jørgen Kjeldsen, Chuan Liu, et al. A reference gene catalogue of the pig gut microbiome. *Nature microbiology*, 1(12):1–6, 2016. [PubMed:27643971] [doi:10.1038/nmicrobiol.2016.161].
